# Functional convergence of genomic and transcriptomic genetic architecture underlying sociability in a live-bearing fish

**DOI:** 10.1101/2023.02.13.528353

**Authors:** Alberto Corral-Lopez, Natasha I. Bloch, Wouter van der Bijl, Maria Cortazar-Chinarro, Alexander Szorkovszky, Alexander Kotrschal, Iulia Darolti, Severine D. Buechel, Maksym Romenskyy, Niclas Kolm, Judith E. Mank

## Abstract

The organization and coordination of fish schools provide a valuable model to investigate the genetic architecture of affiliative behaviors and dissect the mechanisms underlying social behaviors and personalities. We used quantitative genetic methods in replicate guppy selection lines that vary in schooling propensity to investigate the genetic basis of sociability phenotypes. Alignment and attraction, the two major traits forming the sociability personality axis in this species, showed heritability estimates at the upper end of the range previously described for social behaviors, with important variation across sexes. Consistent with findings in collective motion patterns, experimental evolution of schooling propensity increases the sociability of female, but not male, guppies when swimming with unfamiliar conspecifics, a highly relevant trait in a species with fission-fusion social systems in natural populations. Our results from PoolSeq and RNASeq convergently identify genes involved in neuron migration and synaptic function, highlighting a crucial role of glutamatergic synaptic function and calcium-dependent signaling processes in the evolution of schooling.

## Introduction

Living in groups, a widespread phenomenon across the animal kingdom, can lead to strikingly complex social behaviors, such as cooperative interactions, subdivision of labor, or collective decision-making^1^. Sociability, the propensity to affiliate with other animals, can also vary across individuals. Sociability represents a fundamental aspect of personality which can influence social interactions and is often subject to strong selective processes^2, 3^. Indeed, intra-specific differences in sociability are widespread^e.g^ ^4, 5^ and often individual genetic variation underlies variability in personality and social behavior phenotypes^6^. However, heritability estimates of social behavior traits are consistent with a complex, polygenic architecture^7^. Human twin and family studies reveal heritability estimates of personality traits generally ≈ 0.40^(reviewed^ ^in^ ^8)^. In non-human animals, a meta-analysis estimated the mean heritability is 0.23 across social behaviors, including personality traits^9^, with heritability of affiliative associations ranging substantially, from 0.11 to 0.51^10–13^.

Despite this complexity, multiple neural and genetic mechanisms underlying social behavior have been identified^14^. Many of the neural structures and neuromodulators (serotonin, dopamine, vasopressin and oxytocin) are highly conserved within the social-decision making network across vertebrates^15^. Moreover, human personality traits associated with social-decision making have been linked to dopaminergic and serotonergic genes^(reviewed in 16)^, and the regulation of these neuromodulators has been connected to neurodevelopmental disorders that affect affiliative behaviors, such as autism spectrum disorder^6, 17, 18^. Studies in non-human organisms likewise point towards a major role of genes involved in the regulation of these neurochemical systems. For instance, mouse knock-out mutants for genes involved in dopaminergic signalling exhibit altered sociability phenotypes^19^, and changes in sociability in three-spined sticklebacks, *Gasterosteus aculeatus*, are predicted by natural variation in the expression of genes within the dopaminergic and stress pathways^20^. However, while specific groups of genes have been identified for a range of affiliative behaviors, we lack a deeper understanding of their role in inter-individual variation and evolutionary processes underlying sociability.

In fish, group-living often leads to spectacular forms of collective behavior, with members of a school coordinating their movements to increase efficiency in foraging, travelling or predator avoidance^1^. The extent to which members of a school coordinate their movements is an integral part of the sociability axis of personality, i.e., how individuals react to the presence or absence of conspecifics excluding aggressive behaviors^21^. We previously showed that schooling behavior has a repeatability of 0.43 at the individual level^22^, and that can increase substantially over just three generations of artificial selection in female guppies, *Poecilia reticulata*, generating a 15% increase in intrinsic schooling propensity compared to controls^22, 23^. Selection was based on a group phenotype, polarization, or the level of alignment between individuals moving together in a group.

Understanding the genetic basis of this schooling phenotype requires linking individual phenotypic differences to genetic variation. In this study, we phenotyped alignment and attraction of 1496 guppies across 195 families (father, mother, three female and three male offspring from our replicate experimental selection lines) to estimate the heritability of these two motion characteristics that previous factor analyses identified to be integral components for the sociability axis of personality in this species^21^. Because many social interaction patterns in guppies have sex differences^24–26^, and because our selection was performed solely on females, we are able to examine cross-sex genetic correlations in this ecologically relevant behavioral trait. Genomic and transcriptomic data from these lines reveal convergence in the genetic architecture of sociability, highlighting a series of genes with well-defined roles in neurodevelopmental processes. Our results provide a robust agreement across experiments about the genetic regulation of neural processes in decision-making and motor control regions of the brain, and its importance for variation of personality within individuals of this species.

## Results

### Social interactions with unfamiliar individuals and the heritability of sociability in guppies

We first determined whether experimental evolution for higher schooling propensity affected social interactions with unfamiliar conspecifics. For this, we assessed sociability in 740 females and 746 males from multiple families of three replicate lines artificially selected for higher polarization (polarization-selected lines hereon) and three replicate control lines exposed to a group of non-kin unfamiliar conspecifics in an open field test. Specifically, we quantified their alignment and nearest neighbor distance (attraction), two measurements of collective motion characteristics that are demonstrated to capture the most biologically relevant aspects of the sociability axis of personality in this species^21^.

Female guppies from polarization-selected lines presented higher alignment and higher attraction to an unfamiliar group compared to control lines (Linear Mixed Model for alignment, LMM_alignment_: line: t = 2.27; df = 9.68; p = 0.047; LMM_attraction_: line: t = -2.34; df = 9.41; p = 0.043; Fig. 1A; Supplementary Table 1). No differences were observed in these traits between polarization-selected and control males (LMM_alignment_: line: t = -1.38; df = 9.56; p = 0.20; LMM_attraction_: line: t = 0.88; df = 9.26; p = 0.40; Fig. 1A; Supplementary Table 2). Our analyses showed an effect of sex in alignment, with females exhibiting approximately 8% higher alignment than males (LMM_alignment_: sex: t = -3.02; df = 690.08; p = 0.003), but no difference between sexes in attraction to a group of unfamiliar conspecifics (LMM_attraction_: sex: t = 0.51; df = 447.05; p = 0.61; Fig. 1A; Supplementary Table 1). There were some differences in sociability between the parental and offspring generation tested in our experiment, with higher alignment to group average direction and lower distances to nearest neighbor observed in offspring (LMM_alignment_: generation: t = -10.13; df = 1141.24; p < 0.001; LMM_attraction_: generation: t = 11.29; df = 992.16; p < 0.001; Fig. 1A; Supplementary Table 1). Differences in body size between age classes in guppies may explain these results (see Supplementary Table 3), as the time restrictions involved in testing large numbers of fish required that we assessed individuals from the parental and offspring generations at different ages. However, these differences are unlikely to create large biases in our heritability estimates given that we tested all fish after sexual maturation and that polarization-selected and control fish were of similar age within parents tested (9 months old) and within offspring tested (5 months-old). In addition, the difference in means between generations is accounted for in our statistical models (see Methods).

**Figure 1.**
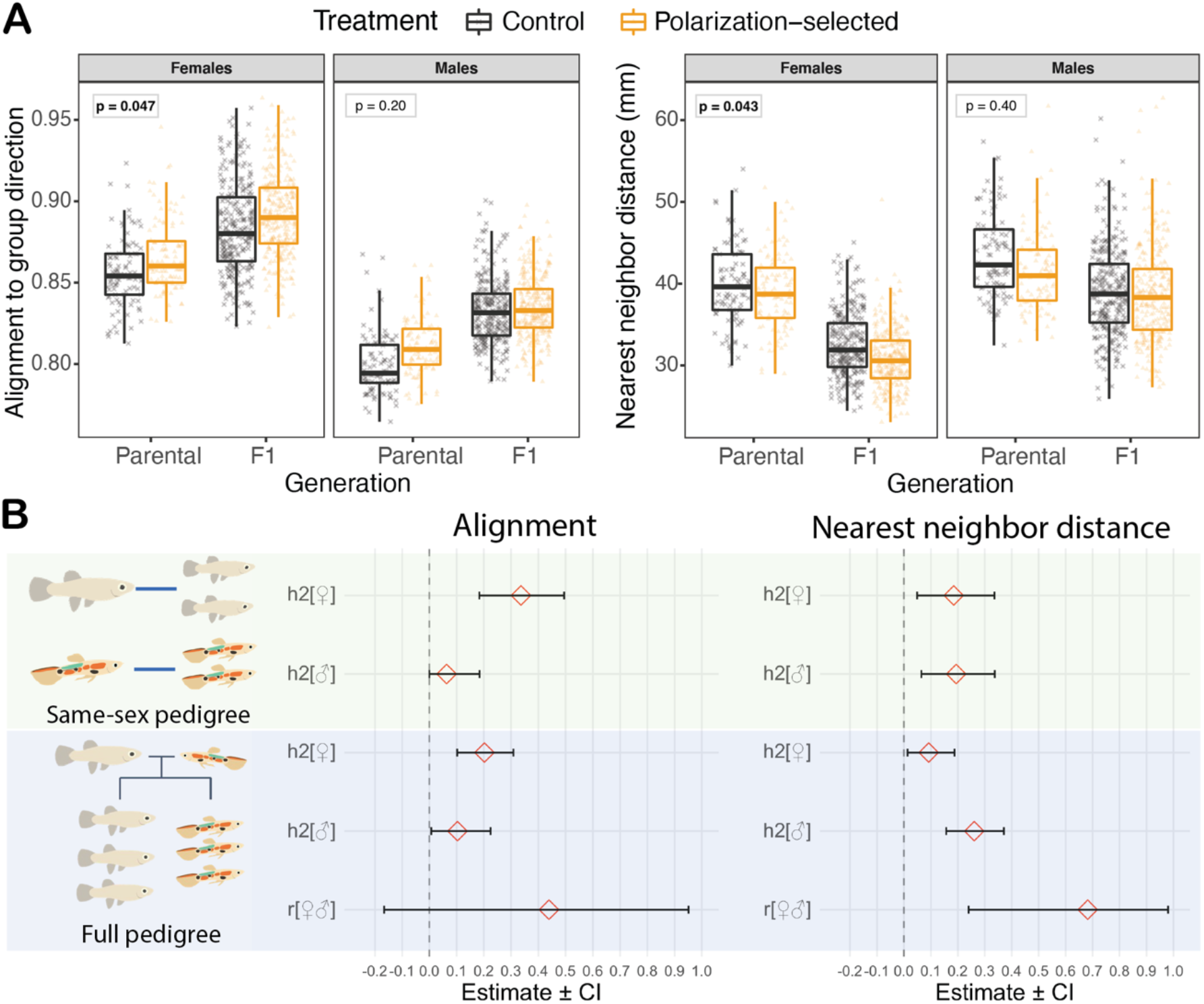
Heritability of sociability in guppies. **(A)** Female, but not male, guppies from polarization-selected lines (orange) presented higher alignment to the group direction (left panel) and shorter distances to their nearest neighbor (higher alignment; right panel) than guppies from control lines (gray) when swimming with unfamiliar same-sex conspecifics (see Supplementary Table 1-2). For all boxplots, horizontal lines indicate medians, boxes indicate the interquartile range, and whiskers indicate all points within 1.5 times the interquartile range. Boxes in top left position of each facet indicate p-values (p < 0.05 in bold) for statistical contrasts by sex in Linear Mixed Model comparing alignment and attraction between selection line treatments (see Supplementary Tables 1-2). **(B)** Animal models using same-sex pedigrees and full pedigrees with alignment and attraction (nearest neighbor distance) phenotypes in 195 families of polarization-selected and control guppy lines indicated a moderate heritability in female guppies of both biologically-relevant aspects of sociability measured, alignment (left) and attraction (right). In males, we found moderate heritability in attraction, but credible intervals in alignment estimates overlapped with 0 suggesting low heritability of this sociability aspect. Our full pedigree animal models provided large credible intervals for male-female correlations in sociability, with estimates overlapping 0 in alignment, but a positive correlation between sexes in attraction (see Supplementary Tables 4-5.

To assess the heritability of sociability in this species, we fitted animal models with alignment and attraction phenotypes quantified from these 1486 individuals, comprised of parents, three male and three female offspring for 195 families (99 polarization-selected and 96 control-families). Given known differences in guppies between the sexes in social interaction patterns^24–26^, we estimated heritability with animal models that only included relationships with same sex individuals (same-sex pedigree) or that include relationship with individuals from both sexes (full pedigree).

Using same-sex pedigree animal models, attraction heritability was similar in females (h^2^ _attraction_, estimate [95% credible interval] = 0.18 [0.05, 0.34]; Fig. 1B) and males (h^2^ _attraction_ = 0.19 [0.06, 0.34]; Fig. 1B), however alignment heritability was far higher in females (h^2^ _alignment_ = 0.34 [0.18, 0.49]; Fig 1B) than males (h^2^ _alignment_ = 0.06 [0.00, 0.18]; Fig. 1B). Full-pedigree models indicated lower heritability estimates than those obtained from same-sex pedigree models (Supplementary Table 5; Fig 1B), except for the heritability estimate of attraction in males (h^2^ _attraction_ = 0.26 [0.16, 0.37]; Fig 1B). Finally, animal models indicated a positive female-male genetic correlation in attraction (r_fm_ attraction: 0.68 [0.23, 0.98]; Fig. 1B), although the magnitude of this correlation contained large credible intervals. For alignment, credible intervals for r_fm_ are also wide and span zero (r_fm_ alignment: 0.44 [-0.17, 0.95]), and we can only conclude that the cross-sex genetic correlation is not strongly negative (Supplementary Table 5; Fig 1B).

### Genetic basis of sociability in guppies

Our quantitative genetic analyses of alignment and attraction suggest an important genetic influence in sociability phenotypes of guppies. As such, we sequenced DNA pools (Pool-seq) to identify genome-wide differences in allele frequencies between polarization-selected female guppies that presented high sociability and polarization-selected females that presented low sociability. Specifically, we focused on measurements obtained from females in analyses of alignment to an unfamiliar group. For this, we pooled the DNA from mothers whose family (normalized mother and daughters alignment score; see methods) were in the top 25% and the bottom 25% quartiles from each of the three replicated polarization-selected lines (six total pooled samples with 7 individuals each; Supplementary Figure 1).

DNA reads were aligned to the guppy reference genome (Guppy_female_1.0 + MT, RefSeq accession: GCA_000633615.2) to compare genome-wide allele frequency differences between high and low sociability guppies. We ran two independent analyses with these aligned sequences. For our first analysis, we merged sequences from the three replicates with high sociability pooled samples and sequences from three replicates with low sociability pooled samples. We filtered merged sequences to 3,004,974 SNPs (see methods) and performed a Fisher’s exact test in Popoolation2^27^ to identify SNPs that significantly differed in their allele frequencies between guppies with high and low sociability. Using this methodology, we identified 819 SNPs associated with our sociability phenotype (Fisher’s exact test, p < 10^-8^; Fig 2A). SNPs over this standard genome-wide significance threshold^28^ were mostly found in single physically unlinked positions across the genome, consistent with a polygenic architecture of the trait.

**Figure 2.**
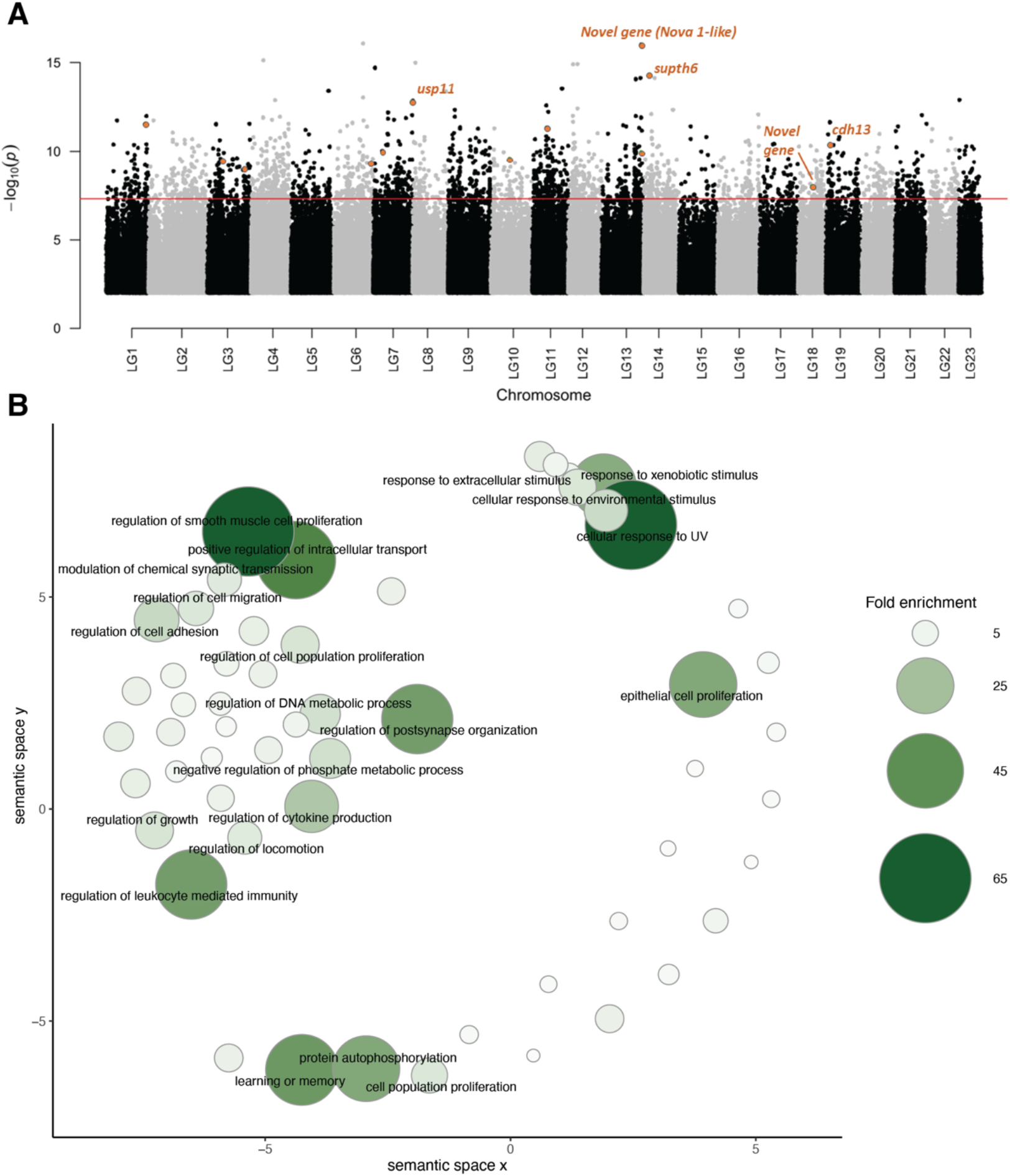
Genetic basis of sociability in the guppy. **(A)** Manhattan plot of -log10(p) values across the guppy genome resulting from a Fisher’s exact test comparing allele frequency differences between high and low sociability female guppies. We merged pooled-DNA sequences of three independent replicates and found 819 SNPs significantly different (above genome-wide threshold highlighted in red), while a highly stringent analyses of consistent allele frequency differences across our three independent replicates (cmh-test; see methods) identified 13 SNPs (five of them within genes) associated with sociability in the species (gene names and SNP location in the genome highlighted in orange). SNPs with -log10(p) values < 2 are omitted. **(B)** Clustering of statistically significant overrepresented Gene Ontology annotations for biological processes associated with differences between high and low sociability in guppies. Point size and color provide information on fold enrichment value from the statistical overrepresentation test performed in PANTHER^84^ (see methods). Terms with fold enrichment lower than 8 are represented but not described in text. Axes have no intrinsic meaning and are based on multidimensional scaling which clustered terms based on semantic similarities^74^.

Out of these 819 significantly different SNPs, 421 were located within genes or gene promoter regions of the guppy genome and were used for further functional characterization in association with Gene Ontology (GO) annotations (273 unique genes). We clustered GO terms based on semantic similarities and found significant overrepresentation of biological process terms related to learning and memory, synaptic functioning, response to stimulus, locomotion and growth (Fig. 2B). We likewise found significant overrepresentation of cadherin and calcium-dependent protein binding annotations (molecular components terms; Supplementary Figure 2) and glutamatergic synapse annotations (cellular components terms, Supplementary Figure 3).

Second, we looked for consistent differences in allele frequencies between high and low sociability pooled samples in our three replicates by performing the Cochran-Mantel-Haenszel test (cmh-test) in Popoolation2^27^. Convergent changes in allele frequency likely represent selected sites, and are less likely the result of genetic drift in any one line. This stringent analysis identified 13 SNPs from ten different chromosomes with consistent significant differences in allele frequencies across the three replicates (cmh-test p-value <0.01 with false discovery rate correction). Five of these SNPs are located within known coding sequence of the guppy genome, of which three are within well-characterized genes in zebrafish and human homologs with important roles for cognitive function – ubiquitin specific peptidase 11 (*usp11*), *supt6* histone chaperone and transcription elongation factor homolog (*supt6h*), and cadherin 13 (*cdh13*; Fig 2A, Table 1). The other two are classified as novel genes, one of them is matched to an RNA-binding protein *Nova-1-like* gene, similarly associated with motor function and changes in synaptic function (Fig 2A, Table 1).

**Table 1.** Characterization of genes associated with sociability in guppies.

| SNP location | Gene ID<br>Ensembl | Gene name | Described cognitive function of<br>homologs | References |
| --- | --- | --- | --- | --- |
| **Chr 7:** 30748139 | 00000004543 | *usp11* | Control of cortical neurogenesis and neuronal migration.<br>Mutations of the gene have been associated with neurological disorders. | Chiang et al, 2021<sup>44</sup><br>Karaca et al. 2015<sup>91</sup> |
| **Chr 13:** 31383940 | 00000018946 | *Novel gene (RNA-binding protein Nova-1-like)* | Neuronal RNA binding protein associated with motor function. | Buckanovich and Darnell 1997<sup>46</sup> |
| **Chr 14:** 4286109 | 00000009725 | *supt6h* | Substrate of mTOR, a signaling pathway associated with brain function and neurodegenerative disorders. | Lipton and Sahin 2015<sup>42</sup><br>Zhao et al. 2015<sup>43</sup> |
| **Chr 18:** 4286109 | 00000014318 | *Novel gene* | - | - |
| **Chr 19:** 3032099 | 00000019822 | *cdh13* | Modulation of brain activity through GABAergic function.<br>Organization of neuronal circuits. | Rivero et al 2015<sup>47</sup><br>Takeichi 2006<sup>92</sup> |

### Neurogenomic response of schooling in guppies

We used transcriptome sequencing to determine differences in gene expression in multiple brain regions of polarization-selected and control females in response to two different social contexts, swimming alone (the “Alone” condition) or schooling in a group (groups of eight unfamiliar females; the “Group” condition). We focused on three separate brain tissues that control distinct functions. The *optic tectum* is involved in sensory processing of visual signals. The *telencephalon* is implicated in decision making. The *midbrain* is associated with motor function in response to auditory and visual stimuli^29, 30^. Together, these three brain tissues contain the main components of the social brain network in fish^31, 32^.

#### Differential expression analyses

We identified genes differentially expressed between lines under each treatment condition and in each brain region separately, to determine the neurogenomic response triggered by schooling in both lines. Gene expression analyses indicated very little overlap in differentially expressed (DE) genes between polarization-selected and control lines (Fig. 3, Supplementary Dataset 1). Specifically, we found that only Adipocyte enhancer-binding protein 2 gene (*AEBP2;* involved in adipocyte differentiation) in the midbrain and an unknown gene in the optic tectum were differentially expressed in both polarization-selected and control lines. Such little overlap suggests that females from different selection lines are activating different transcriptional cascades and biological pathways in response to social context. In polarization-selected lines we found an order of magnitude fewer DE genes in the optic tectum than in the other brain components (n=21 for optic tectum, n=158 for telencephalon, n=109 for midbrain, each p_adj_ < 0.05). Moreover, in the telencephalon and midbrain, DE genes between Alone and Group treatment in the polarization-selected lines were enriched for GO annotations associated with cognition, memory, learning and social behavior (Supplementary Dataset 1). We found enrichment for these annotation terms for DE genes expressed in the optic tectum, but not in the midbrain or telencephalon, of control lines.

**Figure 3.**
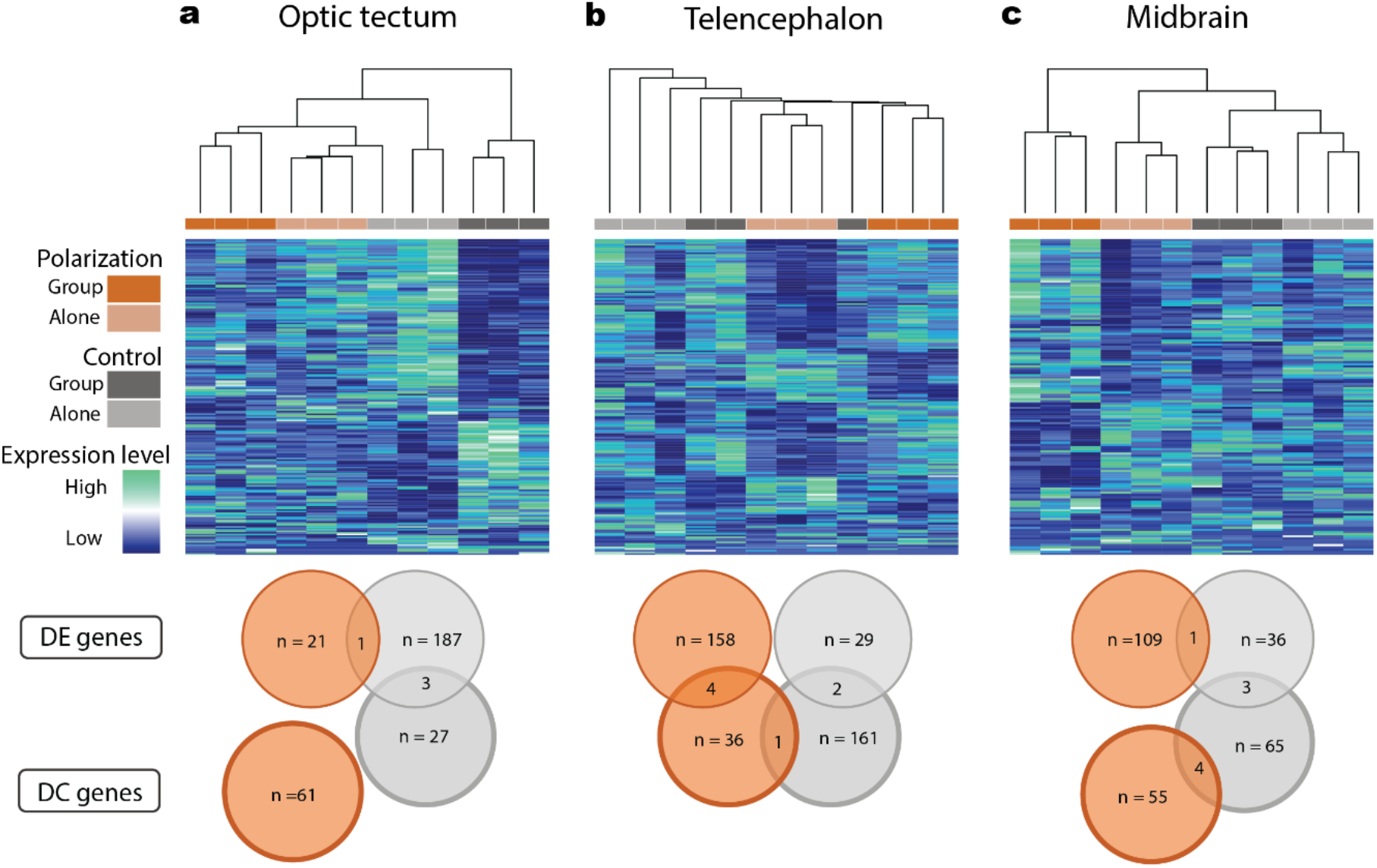
Neurogenomic response of schooling in guppies. Hierarchical clustering and relative expression level for all differentially expressed genes between Alone and Group treatments in the (a) optic tectum, (b) the telencephalon and (c) the midbrain. Differentially expressed genes were identified separately in polarization and control line samples. Clustering, based on Euclidian distance, represents transcriptional similarity across all samples, with bootstrap values (1000 replicates) shown at nodes. Venn diagrams summarize the total number of differentially expressed (DE) genes and differentialy coexpressed (DC) gene pairs in each tissue for polarization-selected and control lines.

Hierarchical clustering analyses of DE genes showed that females from polarization-selected lines in the Group condition clustered uniquely from polarization-selected females in the Alone condition (Fig. 3, Supplementary Table 6). Similarly, females from polarization-selected lines in the Group condition clustered uniquely from females from control lines under the Group and Alone conditions in both the telencephalon and the midbrain, suggesting a unique response in the regions of the brain associated with behavior to social exposure. This was not observed in samples from the optic tectum, suggesting that visual processing of social treatments did not differ between polarization-selected and control females. Hierarchical clustering analyses using all expressed genes clustered samples by selection line rather than by social context condition (Supplementary Fig. 4), suggesting that social context affects only a targeted subset of the overall transcriptome rather than the majority of genes.

#### Differential coexpression analyses

We used systems biology methods designed to compare the coexpression networks between conditions to identify genes that change in the way they are connected to other genes within the coexpression network across conditions, independent of whether they are differentially expressed^29–31^. Specifically, we used BFDCA^30^ to identify differential coexpressed (DC) gene pairs under the Group and Alone conditions (i.e. pairs of genes that significantly change in correlation between the two social contexts for each line^30, 32^). Similar to findings in DE analyses, we found little overlap in the genes forming DC gene pairs between comparisons implemented for control and polarization-selected lines (see Supplementary Tables 7-8, Fig. 3). Together, our results suggest that polarization-selected lines were activating different biological pathways than control lines to modulate coordinated movement.

We additionally found a group of genes that are both differentially expressed (DE) and differentially coexpressed (DC) in the same tissue and line, suggesting they might play an important role mediating coordinated movement (Supplementary Tables 8-9). Specifically, in the telencephalon, we identified 4 genes that are both DE and DC in polarization-selected lines: *LRRC24*, *PTPRS, KHDR2, and PP2BA* (Supplementary Table 9). These genes are part of the Calcineurin and the Wnt/Oxytocin signalling pathways, known to be involved in modulating social behavior, learning and memory^33–35^. Enrichment tests confirm the functional relevance of the DC gene pairs identified, revealing an overrepresentation of genes associated with the glutamatergic synapse, as well as visual transduction among DC gene pairs in multiple comparisons (Supplementary Tables 10-11).

### Functional characterization of genes of interest across experiments

We combined the information from our genomic and transcriptomic analyses on polarization-selected and control lines to obtain an intersected delimitation of the gene functions that our analyses highlighted as important in the development and expression of social interactions with conspecifics. Specifically, we used functional analyses in the set of genes with differentiated SNPs between merged sequences of the three replicates with high and low sociability (273 unique genes) as reference, and compared it to functional analyses of genes differentially expressed in three different brain tissues of females following exposure to multiple social conditions. We found a concordance of 79% in the combination of biological processes (BP), cellular components (CC) and molecular functions (MF) GO terms enriched following analyses of differentially expressed genes in the telencephalon (n =158). This value represented a 1.7-fold increase in the concordance of terms in relation to mean values obtained from corresponding enrichment analyses of 1000 random sets of 158 genes (see Methods; mean concordance ± [CIs]: 45% [43,47]). We likewise found concordances of 64% for differentially expressed genes in the midbrain (n = 109), and of 4.5% for differentially expressed genes in the optic tectum (n = 21). These represented 2.1-fold and 1.1-fold increases in relation to analyses with 1000 random sets of 109 and 21 genes in midbrain and optic tectum respectively (mean concordance midbrain: 30% [28,31]; mean concordance telencephalon: 3.8% [3.6,4.2]). We summarized and visualized GO terms enrichment lists across experiments and tissues sampled using REVIGO^36^. We found a strong overlap between enrichment of GO biological process terms associated with learning and memory, synaptic processes, neuron projection and cell growth, mostly constrained to the telencephalon and midbrain regions (Fig. 4). We found similar patterns in relation to cellular component GO terms, with strong overlap in neuronal components, in particular with high enrichment of terms associated with glutamatergic synapse. Visualization of GO terms associated with molecular functions suggests a major role of genes with protein binding function across experiments, including a role of cadherin-binding related genes in the midbrain (Fig. 4).

**Figure 4.**
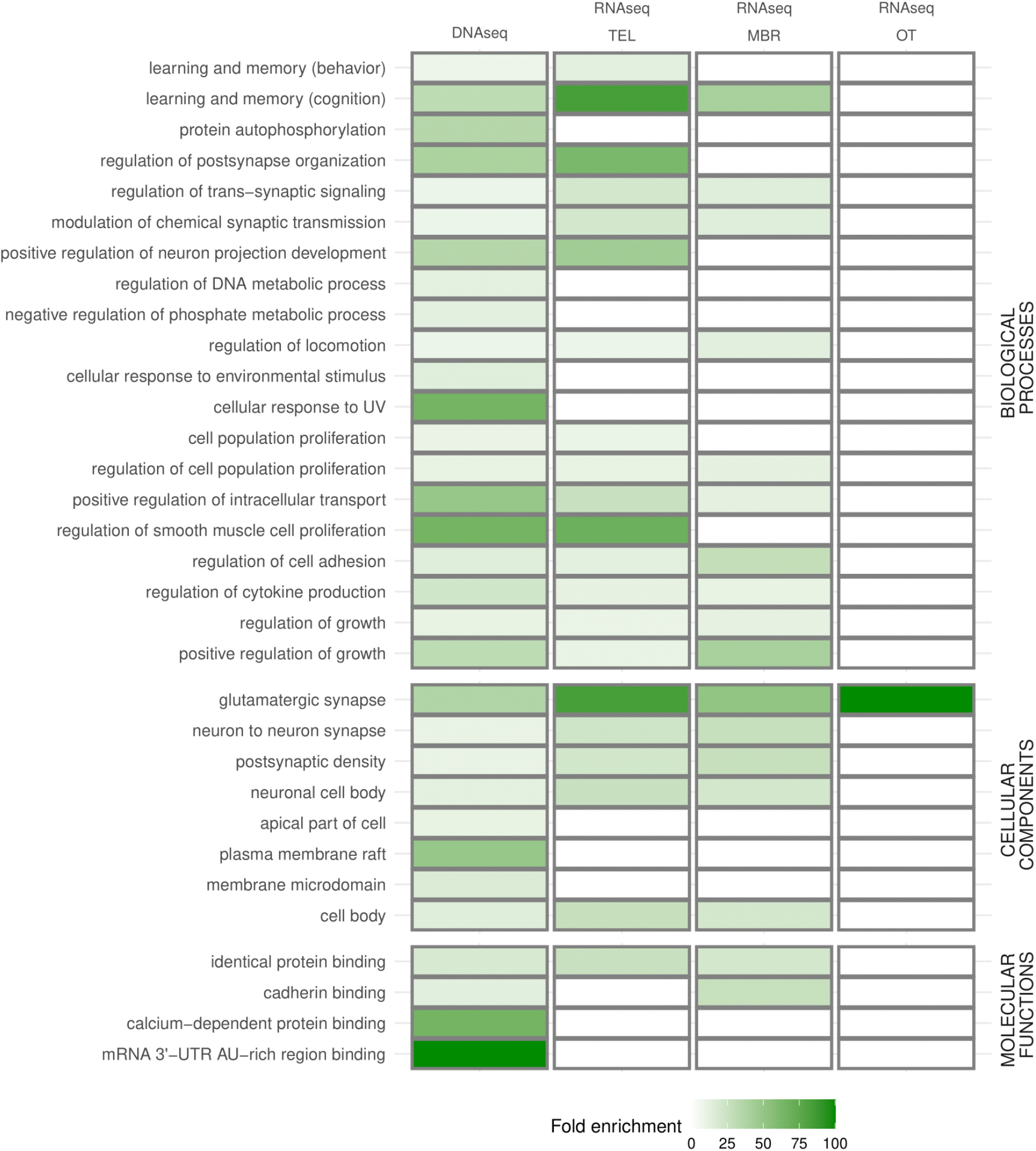
Functional characterization of genes of interest across experiments. Visualization of functional overlap based on Gene Ontology annotations between genes of interest highlighted in strongly differentiated experimental setups evaluating social interactions of female guppies following experimental evolution for higher polarization: 1-genomic analyses of DNA comparing Pool-seq of high and low sociability female guppies (Left Column); 2-transcriptomic analyses evaluating differentially expressed genes in key brain regions of polarization-selected lines of female guppies exposed to two different social contexts, swimming alone or with a group of conspecifics (TEL: telencephalon, MBR: midbrain, OT: optic tectum; Columns 2-4). Shades of green indicate fold enrichment from our statistical overrepresentation tests performed to gene lists obtained from each experiment.

## Discussion

We used behavioral phenotyping across guppy families, in conjunction with PoolSeq and RNASeq to identify the genetic architecture of coordinated motion. Our broad range of analyses, spanning genomes, transcriptomes, and phenotypes, provides an exceptional evaluation of the molecular mechanisms underlying sociability in this fish. Our work suggests that genes and gene networks involved in social decision making through neuron migration and synaptic function are key in the evolution of schooling, highlighting a crucial role of glutamatergic synaptic function and calcium-dependent signaling processes.

Our pedigree-based phenotyping analyses of 195 guppy families from polarization-selected and control lines indicate moderate levels of heritability (Alignment: 0.06 - 0.34; Attraction: 0.09 - 0.26), with pronounced sex differences in full pedigree models (Alignment_female-male_: 0.10 ± 0.05; Attraction_female-male_: -0.17 ± 0.05), in estimates for key behavioral traits forming the sociability axis in this species. Our heritability estimates are similar to estimates for affiliative social behavior traits in primates, ungulates and rodents^10–13, 37^, and to overall estimates of heritability in personality traits across human and non-human animals^8, 38^. Given the importance of social behavior in a range of survival and fitness components in natural systems^1, 39, 40^, our results suggest that complex genetic architectures can respond quickly to strong evolutionary pressures, even when only one sex is subject to selection^22^, and that our lab population contained significant amounts of standing genetic variation for these traits prior to selection.

The complex genetic architecture makes it difficult to precisely characterize cross sex genetic effects in our study. We nonetheless observed a positive cross sex genetic correlation in attraction (0.68, CI: 0.25 - 0.98), suggesting similarities between males and females in the genetic architecture of this trait. This result is concordant with a study focused on the bold-shy continuum aspect of personality establishing that sex differences in risk-taking behaviors are weak and likely lack sex-specific genetic architecture in this species^41^. Yet, sex differences in heritability estimates of alignment (♀h^2^ _alignment_ = 0.34 [0.18, 0.49]; ♂h^2^ _alignment_ = 0.06 [0.00, 0.18]) suggest that it is important to account for sex-specific additive genetic variance when inferring the evolvability of personality traits. Additionally, the low cross-sex heritability we observe in these latter traits is particularly interesting, and suggests that selection in one sex for a complex trait need not result in a correlated response in the other sex. Overall, this indicates significant sex-specific genetic variation for sex-specific behaviours, and that sexually dimorphic behaviors need not require decoupling of male and female genetic architecture when sufficient sex-specific genetic variation is present.

We next mapped the genomic and transcriptomic basis of phenotypic differences in polarization in female guppies. Our GWAS experimental design was designed to compare individuals with high- and low-sociability phenotypes from within polarization-selected lines, rather than between polarization-selected and control lines. This may have resulted in compressed phenotypic spread, but carries the important advantage of reducing the incidence of SNP frequencies that vary across alternative selection lines due to drift. As such, our design is conservative. In our most stringent PoolSeq analysis, we identified SNPs in four genes that consistently differed across all three replicate lines, which have previously been associated with cognition and functions relevant to social behavior. The supt6 histone chaperone and transcription elongation factor homolog (*supt6h*) is important in the positive regulation of transcriptional elongation and a substrate of mTOR, a signalling pathway with a role in cognitive function^42, 43^. Ubiquitin specific peptidase 11 (*usp11*) homolog in humans has a critical function in the development of neural cortex, and knock-out studies in mice show that the locus protects females from cognitive impairment^44, 45^. Similarly, the novel RNA-binding protein *Nova-1-like* gene is associated with a neuron-specific nuclear RNA binding protein in humans and regulates brain-specific splicing related to synaptic function^46^.

Finally, our PoolSeq analysis identified Cadherin 13 (*cdh13*), the human homolog of which has a crucial role in GABAergic function^47^, with involvement in neural growth and axonal guidance during early development^48, 49^. Moreover, deficit of this gene has a major impact in neurodevelopmental disorders including attention-deficit/hyperactivity disorder and autism spectrum disorder^50^. Indeed, *cdh13* knock-out mice display delayed acquisition of learning tasks and a decreased latency in sociability^51^. Interestingly, repeated selection of genes involved in cadherin-signalling pathways^52^ has been shown in guppy populations experiencing different predation pressures. Together, natural selection imposed by differences in predation across these populations^53–55^, and our combined findings in the genomic background of guppies suggest strong selective pressures for cadherin-signalling genes due to their modulation of affiliative behaviors.

Our expression results revealed differences in regulation in genes associated with learning, behavior, and neural function, mainly in the telencephalon and midbrain, in comparisons of polarization-selected and control lines in different social contexts. Overall, this suggests that the regulation of highly demanding cognitive processes via synaptic function underlies variation in sociability. While the integration of visual signals is central in fish schools^56^, our results suggest that higher order cognitive processes are the basis of variation in social affinity. Indeed, the differences in alignment and attraction observed when swimming with unfamiliar conspecifics are arguably highly cognitively demanding, as within a collective motion context, the tendency to copy the directional movements of other individuals implies a direct trade-off between personal goal-oriented behaviors and the benefits of social conformity^57, 58^. Together, our study of transcriptomic profiles of schooling fish suggests that the regulation of affiliative behaviors in this species is driven by an intricately linked social decision-making network in the brain^59^, with strong links to functional groups governing social behaviors and personality across species. More broadly, our results offer insight into important questions about the evolution of behaviour, and other traits with complex genetic architecture. First, our results of large-scale expression differences among selection lines are consistent with recent discussions of the role of gene regulatory networks in co-ordinating large numbers of genes associated with behaviours^60^. It is highly likely that the genes with convergent expression changes in the selection lines are controlled via a modular regulatory architecture, as evidenced by our coexpression network analysis (Supplementary Tables 7-8, Fig. 3).

We find a striking concordance in the functionality of genes independently identified in genomic and transcriptomic profiling of strongly differentiated experiments assessing social interactions of polarization-selected female guppies (Fig. 4). The overlap in significantly enriched Gene Ontology terms, including learning, synaptic processes and neuron projection restricted to brain regions associated with decision-making and motor control, strongly reinforces the notion that genetic regulation of these cognitive processes is fundamental for sociability in guppies. Additionally, the functional concordance we observe between the regulatory and protein differences among our selection lines is noteworthy in the context of the discussion of whether structural or regulatory variation is more important in adaptive phenotypes^61, 62^. The overlap in functionality in our genomic and transcriptomic approaches suggests that both are important, with artificial selection for behavior acting on coding and regulatory variation within the same pathway to achieve adaptive phenotypes.

Our results indicate that the regulation of glutamatergic synaptic processes is a particularly promising network for future studies of affiliative behaviors. Interestingly, differential gene expression of glutamate receptor genes has been identified to regulate female mating preferences in guppies^63^, and is concordantly identified across species of vertebrates in the regulation of long-term affiliative mating behaviors^64^. Guppies are livebearers, and this has hindered the use of functional genetic tools such as CRISPR on this species. Although not feasible at this time, future functional validation via genetic manipulations of guppies of these pathways would prove extremely interesting.

Our results are also concordant with other comparative transcriptomic studies of behavioral responses towards conspecific territorial intrusions, which identified calcium ion-binding regulation across phylogenetically distant species^65^. Together, the consistency in our findings of specific genes and functional terms associated with calcium-dependent and cadherin binding molecular functions across our experiments suggest these are promising molecular targets for future research exploring the evolution and regulation of sociability and affiliative behaviors.

## Methods

### Study system

To evaluate the genetic architecture of sociability, we performed a series of experiments in guppies following artificial selection on coordinated motion. The laboratory population of guppies used originated from a downstream population of the Quare river in Trinidad, which is subject to high predation levels. The original collection was made in 1998^66^, and the laboratory population has since been kept in several large (>200-litre) tanks of >200 individuals each to avoid inbreeding. The artificial selection procedure is outlined in detail in^22, 23^. In brief, groups of female guppies were subjected to repeated open field tests, and were subsequently sorted based on their median polarization, measured by the degree of alignment exhibited by the individuals within the group when swimming together^22, 23^. For three generations, females from groups with higher polarization were mated with males from those cohorts to generate three lines of guppies that had been selected for high polarization. In parallel, random females were exposed to the same experimental conditions and were mated with unselected males to generate three control lines. Analysis of the third generation of polarization-selection revealed that, on average, females exhibited a 15% higher level of polarization and a 10% higher level of group cohesiveness compared to control females^22^.

Throughout the selection experiment and the completion of experiments described below all fish were removed from their parental tanks after birth, separated by sex at the first onset of sexual maturation, and afterwards kept in single-sex groups of eight individuals in 7-L tanks containing 2 cm of gravel with continuously aerated water, a biological filter, and plants for environmental enrichment. We allowed for visual contact between the tanks. The laboratory was maintained at 26°C with a 12-h light:12-h dark schedule. Fish were fed a diet of flake food and freshly hatched brine shrimp daily.

### Social interactions with unfamiliar individuals and the heritability of sociability

To investigate the heritability and cross-sex genetic correlations of sociability in the guppy, we measured alignment and attraction with unfamiliar groups of conspecifics in parents and offspring from polarization-selected and control lines. Specifically, using offspring of the F3 generation of selection we bred 35 families for each of the three polarization-selected and for each of the three control lines. From our population of F3 generation offspring (kept in single sex groups prior to the breeding experiments), we used male and female guppies of the same age (approximately 9 months old) and paired them in 3L tanks to generate the parental generation. We collected offspring for the two first clutches of these pairs and we transferred newborn offspring to 3L tanks in groups of three siblings. We separated siblings by sex at the first onset of sexual maturation, and afterwards we kept them in single-sex groups of three individuals until behavioral testing. We phenotyped sociability of a total of 195 guppy families: mother, father and six offspring (three females and three males). Any family for which we did not collect at least three female and three male offspring was disregarded from further behavioral testing. Each of the six selection lines was represented with a minimum of 30 families in our heritability analyses.

#### Behavioral assays

To phenotype sociability in each member of our guppy families, we measured alignment and attraction of 1495 guppies from our breeding experiment. For each fish, we performed an open field assay using white arenas with 55 cm diameter and 3 cm water depth in which our focal fish (guppies from the breeding experiment) interacted with a group of seven same-sex conspecifics. Non-focal guppies used in these assays were from a lab wild-type stock population and of similar age to our focal fish. Prior to the start of the test, focal fish and the seven-fish group were acclimated in the centre of the arena for one minute in separate opaque white 15 cm PVC cylinders. After this acclimation period, we lifted the cylinders and filmed the arena for 10 minutes using a Point Grey Grasshopper 3 camera (FLIR Systems; resolution, 2048 pixels by 2048 pixels; frame rate, 25 Hz). Three weeks prior to assays, we tagged wild-type fish with small black elastomere implants (Northwest Marine Technology Inc.) to allow recognition of wild-type fish after completion of each assay. After completion, we gently euthanized focal fish from the parental generation with an overdose of benzocaine and kept them in ethanol for future genomic analyses. Focal fish from the offspring generation were transferred to group tanks for future experimental use. Groups of seven wild-type fish were transferred to holding tanks and used in a maximum of seven assays with focal fish.

#### Data processing

We tracked the movement of fish groups in the collected video recordings using idTracker^67^ and used fine-grained tracking data to calculate the following variables in Matlab (v2020): i) alignment, the median alignment of the focal fish to the group average direction across all frames in the assay. This was quantified by the total length of the sum of two-unit vectors, one representing the heading of the focal fish, and one representing the heading of the group centroid. Calculations of alignment were only obtained if six out of the eight members of the group presented tracks following the optimization of our tracking protocol in the setup in (^22, 23, 68^); ii) attraction, the median nearest neighbor distance across all frames in the assay; and iii) activity, we obtained the median speed across all group members and across all frames by calculating the first derivatives of the x and y time series, then smoothed using a third-order Savitzky–Golay filter. For all measurements, trials with less than 70% complete tracks (n= 8) were disregarded for further analyses. The proportion of frames used did not differ between polarization-selected and control fish for any comparison across different generations and sexes (Supplementary Figure 5). We calculated these variables for the focal fish and the average for the seven-fish wild-type group. To recover focal fish id in the tracking data we used idPlayer to visualize trials by projecting the raw tracking data onto experimental videos. We followed focal individuals for the first two minutes of the assay and used the stable identity assigned by idTracker in data collection. In trials with less than 85% complete tracks (n= 8), we followed focal individuals for the total duration of the recording to verify the consistency of identity assigned by idTracker. This approach has previously shown strong reliability in individuals that were observed using this protocol for 20-min recordings in the same experimental setup that quantified sexual behavior of guppies in mixed-sex shoals^69^.

#### Statistical analyses

Analyses were conducted using R Statistical Software (v4.1.3)^70^, RStudio (v2023.3.1.446)^71^ and the tidyverse package^72^. We used Linear Mixed Models (LMMs) with alignment and attraction as dependent variables to test for potential differences between polarization-selected and control lines in social interactions with unfamiliar individuals. Selection regime, sex, the interaction between these two factors and generation were included as fixed effects. The average activity of the wild-type group was coded as a covariate, with a random intercept for each replicated selection line, the breeding family, and the number of tests previously performed with the wild-type group as random factors. All models were run using lme4 and lmerTest packages^73, 74^. Model diagnostics showed that residual distributions were roughly normal with no evidence of heteroscedasticity.

To estimate heritability, the degree of phenotypic variation due to genetic inheritance, and cross-sex genetic correlations of alignment and attraction we used Bayesian animal models^75^. Animal models use a matrix of pedigree relationships set as a random effect, to separate phenotypic variance for each response variable into additive genetic variance and the remaining variance. Given strong sex differences in social interactions in guppies, we performed three animal models for each trait: one including the data on the 1495 phenotyped individuals, and two including only the phenotyped females or males. Parameter values were estimated using the brms interface^76, 77^ to the probabilistic programming language Stan^78^. We used normal priors with a mean of 0 and standard deviation (sd) of 3 for fixed effects, and student-t priors with 5 degrees of freedom, a mean of 0 and sd of 5 for random effects. The full pedigree model estimated cross-sex correlations with a Lewandowski-Kurowicka-Joe (LKJ) prior with η = 1, which is uniform over the range −1 to 1. Posterior distributions for full / same-sex pedigree models were obtained using Stan’s no-U-turn HMC with 24 / 16 independent Markov chains of 2500 / 4000 iterations, discarding the first 1500 / 2000 iterations per chain as warm-up and resulting in 24000 / 32000 posterior samples overall. Convergence of the chains and sufficient sampling of posterior distributions were confirmed by a potential-scale-reduction metric (R) below 1.01, and an effective sample size of at least 1000. For each model, posterior samples were summarized based on the Bayesian point estimate (posterior median) and posterior uncertainty intervals by HDIs. We calculated estimates of heritability by taking the ratio of the additive genetic variance to the total phenotypic variance in each independent model (see Supplementary Tables 5-6).

### Genetic basis of sociability in guppies

#### Pooled DNA sequencing

We extracted DNA of muscle tissue from the caudal peduncle of polarization-selected females from the parental generation using Qiagen’s DNAeasy Blood & Tissue kit following standard manufacturer’s protocol with an additional on-column RNase A treatment. We quantified DNA concentration using fluorometry (Qubit; ThermoFisher Scientific). We next pooled samples from the seven females that represented the top and bottom 20% polarization-selected guppy lines whose families presented higher and lower sociability in six final pools at equimolar amounts (Supplementary Figure 1). We achieved a minimum of three μg genomic DNA per pool. We used a Nextera DNA Flex library preparation kit (Illumina) following the manufacturer protocol. The final library containing six pooled samples was sequenced at SciLife Lab, Uppsala (Sweden) in one lane of an Illumina NovaSeq 6000. We obtained on average 31.8 million 150bp read pairs per sample (26.9 million read pairs minimum per sample).

#### Read quality control and trimming

We assessed the quality of reads for each pool using FastQC v.0.11.4 (www.bioinformatics.babraham.ac.uk/projects/fastqc). After verifying initial read quality, reads were trimmed with Trimmomatic v0.35^79^. We filtered adaptor sequences and trimmed reads if the sliding window average Phred score over four bases was <15 or if the leading/trailing bases had a Phred score <4, removing reads post filtering if either read pair was <50 bases in length. Quality was verified after trimming with FastQC.

#### Genome-wide allele frequency analysis

Reads were mapped using default settings to the guppy reference genome assembly (Guppy_female_1.0 + MT, RefSeq accession: GCA_000633615.2)^80^ with bwa-mem (v0.7.17)^81^. We used Samtools (v. 1.6.0)^82^ to convert sam to bam files, to sort bam files, to remove duplicates, and to make mpileup files. First, to identify SNPs that significantly differed in their allele frequencies between guppies with high and low sociability, we merged sequences from high sociability and low sociability pools and used Popoolation2^27^ to create a synchronized file with allele frequencies for high and low sociability (mpileup2sync.pl –min-qual 20), compute allele frequency differences (mpileup2sync.pl --min-count 6 --min-coverage 25 -- max-coverage 200), calculate Fst for every SNP (fst-sliding.pl) and perform a Fisher’s exact test (fisher-test.pl). Second, we similarly used Popoolation2 to detect consistent changes in allele frequencies of sociability pooled samples for our three replicated artificial selection lines. For this, we created one sync file per replicate (mpileup2sync.pl –min-qual 20), and perform a Cochran-Mantel-Haenszel (CMH) test (cmh-test.pl --min-count 18 --min-coverage 25 --max-coverage 200). Using package qqman^83^ in R (v 4.1.3)^70^ we made Manhattan plots for each chromosome by plotting the negative log10-transformed p-values of exact fisher and CMH tests as a function of chromosome position.

#### Significance tests and functional analyses

We determined SNPs that were significantly different between high and low sociability merged pools in Fisher’s exact tests using the traditional genome-wide significance threshold [–log10(p) > 8]^28^. We next used custom scripts to identify the overlap between position of these SNPs and genes present in the guppy reference annotated genome^80^, and to find homologous genes of these set in medaka (*Oryzias latipes*). We further used this set of unique genes (n = 160) to determine associated GO terms between our merged pools. For this, we performed enrichment tests in PANTHER^84^, as implemented in the GO Ontology Consortium (http://www.geneontology.org/). To test for enrichments of GO terms we performed one-tail Fisher’s exact tests with a Bonferroni corrected p-value threshold of p < 0.05 using a full list of medaka genes orthologous to guppy genes as background. We used Revigo (http://revigo.irb.hr)^36^ to find and visualize representative subsets of terms based on semantic similarity measurements for our enriched GO terms related to biological processes, cellular components and molecular functions.

For CMH test results, we determined SNPs that were significantly different between high and low sociability pools based on FDR corrected p-value < 0.01. We used a custom script to identify the overlap between position of these SNPs and genes present in the guppy reference annotated genome^80^.

### Neurogenomic response of schooling in guppies

#### Behavioral assays and tissue collection

Using offspring of the F3 generation (6-months-old), we placed an individual or groups of eight unfamiliar adult control and polarization-selected females in white 55cm arenas. After 30 min females were euthanized by transfer to ice water. After 30 seconds, with aid of a Leica S4E microscope, we removed the top of the skull and after cutting transversally posterior of the optic tectum and anterior of the cerebellum, and horizontally through the optic chiasm, removed the brain from the skull and placed it into ice water. We severed the *telencephalon* from the rest of the brain between the ventral telencephalon and thalamus at the ‘commissura anterioris’, including both the pallium and subpallium regions. Then we cut the laminated cup-like structures of the *optic tectum*. The remaining part of the brain was the *midbrain*. Dissections took under 2 minutes and tissue samples were immediately preserved in RNAlater (Ambion) at 4°C for 24 hours and then at -20°C until RNA extraction.

#### RNA extraction and sequencing

For each treatment, we pooled tissue from ten individuals into two non-overlapping pools of five for each replicate line. We used this strategy to reduce noise in transcript expression data during sample normalization procedures potentially caused by outliers during behavioral experiments, while maintaining each replicate as a comparable unit. Our experimental design represents a total of 120 individual females, constituting six pools per treatment, per selection regime for a total of 24 pools per tissue. Each sample pool was homogenized, and RNA was extracted using Qiagen’s RNAeasy kits following standard manufacturer’s protocol. Libraries for each sample were prepared and sequenced by the Wellcome Trust Center for Human Genetics at the University of Oxford, UK. All samples were sequenced across nine lanes on an Illumina HiSeq 4000. We obtained on average 33.9 million 75bp read pairs per sample (28.9 million read pairs minimum, 39.8 million maximum).

#### Read quality control and trimming

We assessed the quality of reads for each sample using FastQC v.0.11.4 (www.bioinformatics.babraham.ac.uk/projects/fastqc). After verifying initial read quality, reads were trimmed with Trimmomatic v0.35^79^. We filtered adaptor sequences and trimmed reads if the sliding window average Phred score over four bases was < 15 or if the leading/trailing bases had a Phred score < 3, removing reads post filtering if either read pair was < 33 bases in length. Quality was verified after trimming with FastQC.

#### Differential Expression Analysis

We mapped RNAseq reads against the latest release of the published guppy genome assembly^80^, using the HiSat 2.0.5 – Stringtie v1.3.2 suite^81^. For each individual pool, reads were mapped to the genome and built into transcripts using default parameters. The resulting individual assemblies were then merged into a single, non-redundant assembly using the built-in StringTie-merge function. We filtered the resulting assembly for non-coding RNA using medaka and Amazon molly (*Poecilia formosa*) non-coding RNA sequences as reference in a nucleotide BLAST (Blastn). After eliminating all sequences matching non-coding RNAs, we kept only the longest isoform representative for each transcript for further analysis. Finally, we quantified expression by re-mapping reads to this filtered assembly using RSEM (v1.2.20)^85^.

Lowly expressed genes were removed by filtering transcripts < 2 RPKM (reads per kilobase per million mapped reads), preserving only those transcripts that have expression above this threshold in at least half of the samples for each treatment within a line. After this final filter, a total of 26,140 optic tectum transcripts, 25,100 telencephalon transcripts and 26,514 midbrain transcripts were retained for further analysis. Using sample correlations in combination with MDS plots based on all expressed transcripts, we determined that none of the 72 pools represented outliers and thus, all samples were included in the analysis.

We used DESeq2^86^ to normalize filtered read counts using standard function to identify differentially expressed (DE) genes between the alone and the group treatment in control and polarization-selected lines separately, and then examined the overlap in differentially expressed genes between them. A transcript was considered differentially expressed if it had an FDR corrected p-value < 0.05. As behavior could be modulated by small changes in expression, we did not filter differentially expressed genes based on Log Fold-change in expression between the treatments.

#### Differential Coexpression Analysis

We used Bayes approach for Differential Coexpression Analysis (BFDCA)^30^, in order to identify pairs of genes that have different correlation patterns in two conditions^32, 87, 88^. Here we compared the Alone and Group treatments within each line for each tissue separately, in the same manner as the previously described DE analysis. BFDCA is based on WGCNA and has been shown to be a reliable and accurate method^30^. This untargeted approach to differential coexpression (DC) analysis uses a combined Bayes factor, a ratio of marginal likelihood of the data between the two alternative hypotheses, to evaluate which genes are differentially correlated in two conditions. We controlled for false positives and accounted for multiple testing by integrating a random permutations approach^32^. In short, we created 1000 permuted datasets and considered a DC gene pair significant if the Bayes factor for the actual expression data was larger than the 1% tail of the permutated data Bayes factor distribution.

#### Functional Analyses

To investigate the function of DE genes we performed GO term enrichment tests. To accomplish this, we initially completed the annotation of the reference genome assembly. The transcripts without clear gene names from the reference genome, and the de novo transcripts identified by HiSat were annotated with blastX against the Swissprot non-redundant database. We then determined which GO terms were associated with differentially expressed genes and performed Biological Processes (BP), Cellular Components (CC) and Molecular Functions (MF) enrichment tests in PANTHER^84^. To assess the level of concordance between genes of interest across experiments, we compared the proportion of BP, CC and MF GO terms that were significantly enriched in genomic analyses of sociability implemented in polarization-selected females, and the proportion of BP, CC and MF GO terms enriched in differential expression analyses in brain tissue of polarization-selected females following exposure to Group and Alone experimental conditions. To assess their significance, we compared these values to mean proportions obtained from bootstrap analyses of 1000 random sets of 158 (for comparison with telencephalon), 109 (midbrain) and 21 (optic tectum) genes from our medaka-guppy orthologous gene list. All analyses were based on one-tail Fisher’s exact tests with a Bonferroni corrected p-value threshold of p < 0.05 using medaka genes orthologous to guppy genes as the background. Bootstrap analyses with random sets of genes were automated using rbioapi package^89^ in R (v4.1.3)^70^. We next summarized and visualized GO terms enrichment lists across experiments and tissues using REVIGO^36^ (settings: SimRel semantic similarity measure, 0.5 value). To investigate the function of differentially coexpressed genes, we used g:Profiler^90^ to identify the enriched BP GO terms and pathways that were altered across mating contexts associated with differentially coexpressed gene pairs. We determined over-represented pathways among DC gene pairs in each tissue using the human (Homo sapiens) database in g:Profiler. We chose the human database for its completeness, acknowledging the distant phylogenetic relationship to guppies.

## Acknowledgements

We thank David Sumpter, Kristiaan Pelckmans and James Herbert-Read for important contributions to the conceptualization of the artificial selection procedure. We thank three anonymous reviewers for comments and suggestions to revise a previous version of the manuscript. We are grateful to Jacelyn Shu for the design of guppy graphics for figures. We thank Anna Rennie, Emilio Trejo, and Annika Boussard for help with fish husbandry.

## Author contributions

J.E.M., N.K. and A.C-L. contributed to conceptualization and funding acquisition of the project. N.K., A.K., A.S., and M.R. contributed to the design of the selection procedure and behavioral experiments. A.C-L. conducted research to obtain behavioral data. A.C-L. and W.vdB. performed formal analyses and visualization of behavioral and heritability data. A.C-L., M.C-C. conducted research to obtain genomic data. A.C-L., M.C-C. and I.D. perform formal analyses and visualization of genomic data. N.B. and A.K. conducted research to obtain transcriptomics data. N.B. and. A.C-L. performed formal analyses and visualization of transcriptomics data. A.C-L., J.E.M., and N.K. wrote the original draft. All authors contributed to the final version of the manuscript.

## Data and materials availability

Data and code needed to support the conclusions in the paper are deposited in figshare repository (doi: 10.6084/m9.figshare.23805702). Genomic and transcriptomic data is deposited at NCBI (accession codes PRJNA994132 and PRJNA504011). Additional data related to this paper may be requested from the authors.

## Supplementary Figures for

**Figure S1.**
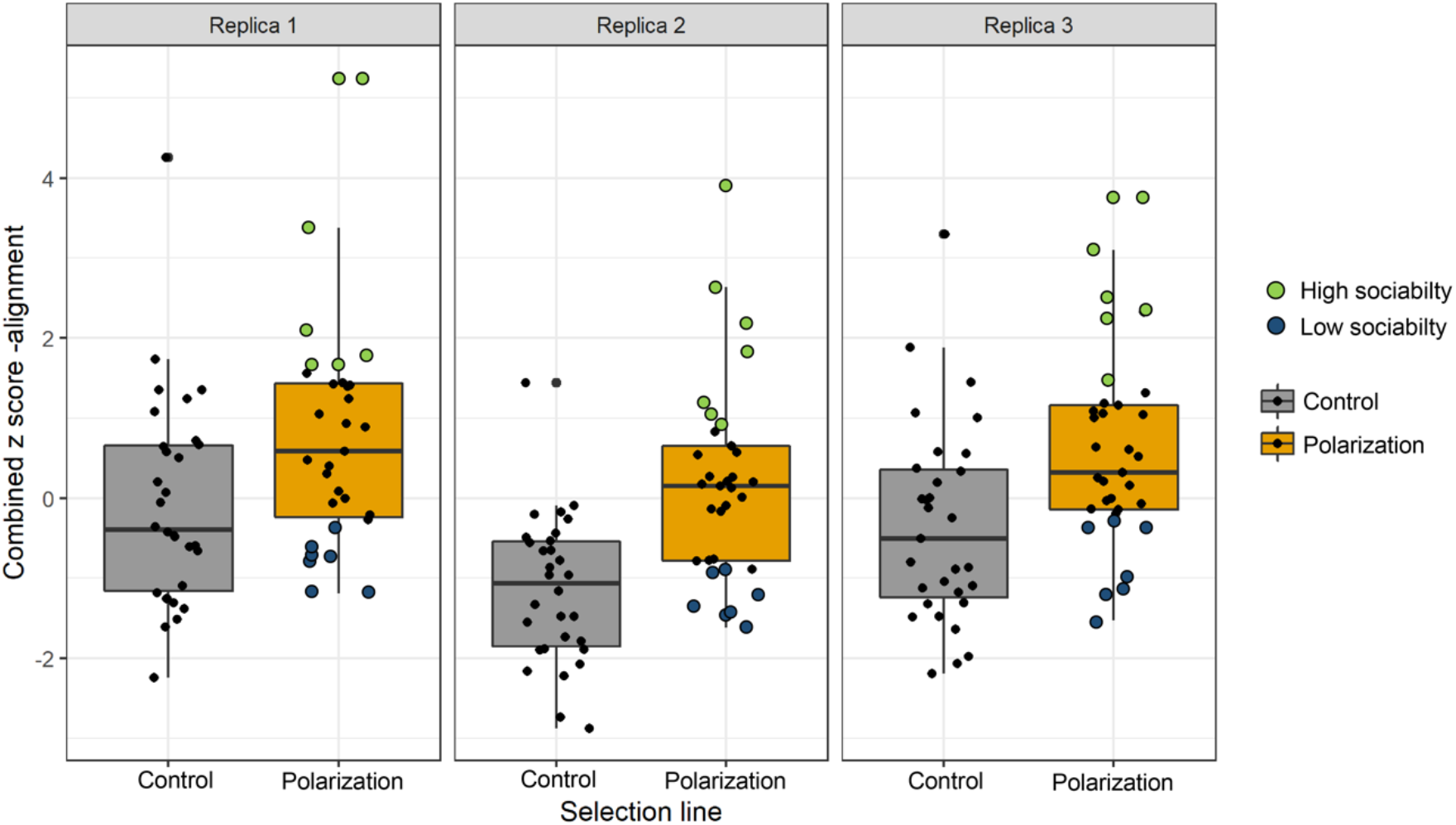
Mean score for alignment to group direction in females for each of the 195 families measured when evaluating the heritability of sociability by means of open field tests in which guppies from three independently replicated polarization-selected (orange) and control (gray) lines were exposed to a group of other seven conspecifics. We obtained combined Z scores by standardizing independently alignment scores obtained in maternal and offspring generations and by adding the score of the mother to the mean score of all female offspring measure in each family. For all boxplots, horizontal lines indicate medians, boxes indicate the interquartile range, and whiskers indicate all points within 1.5 times the interquartile range. Colored circles indicate families that were further used for genomic analyses.

**Figure S2.**
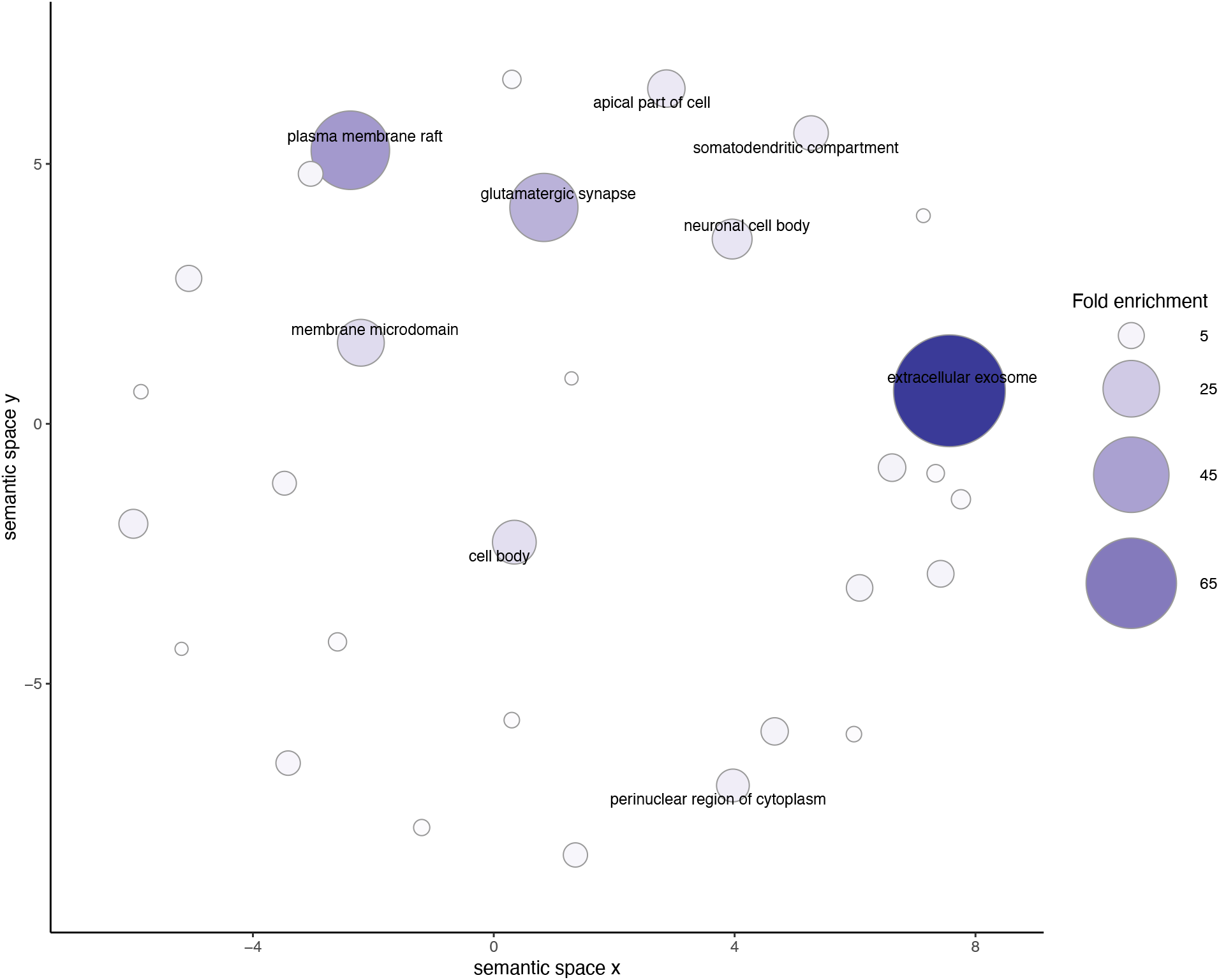
Clustering of statistically significant overrepresented Gene Ontology annotations for cellular components associated to differences between high and low sociability in guppies. Point size and color provide information on fold enrichment value from the statistical overrepresentation test performed in PANTHER^1^. Terms with fold enrichment lower than eight are represented but not described in text. Axes have no intrinsic meaning and are based on multidimensional scaling which cluster terms based on semantic similarities^2^.

**Figure S3.**
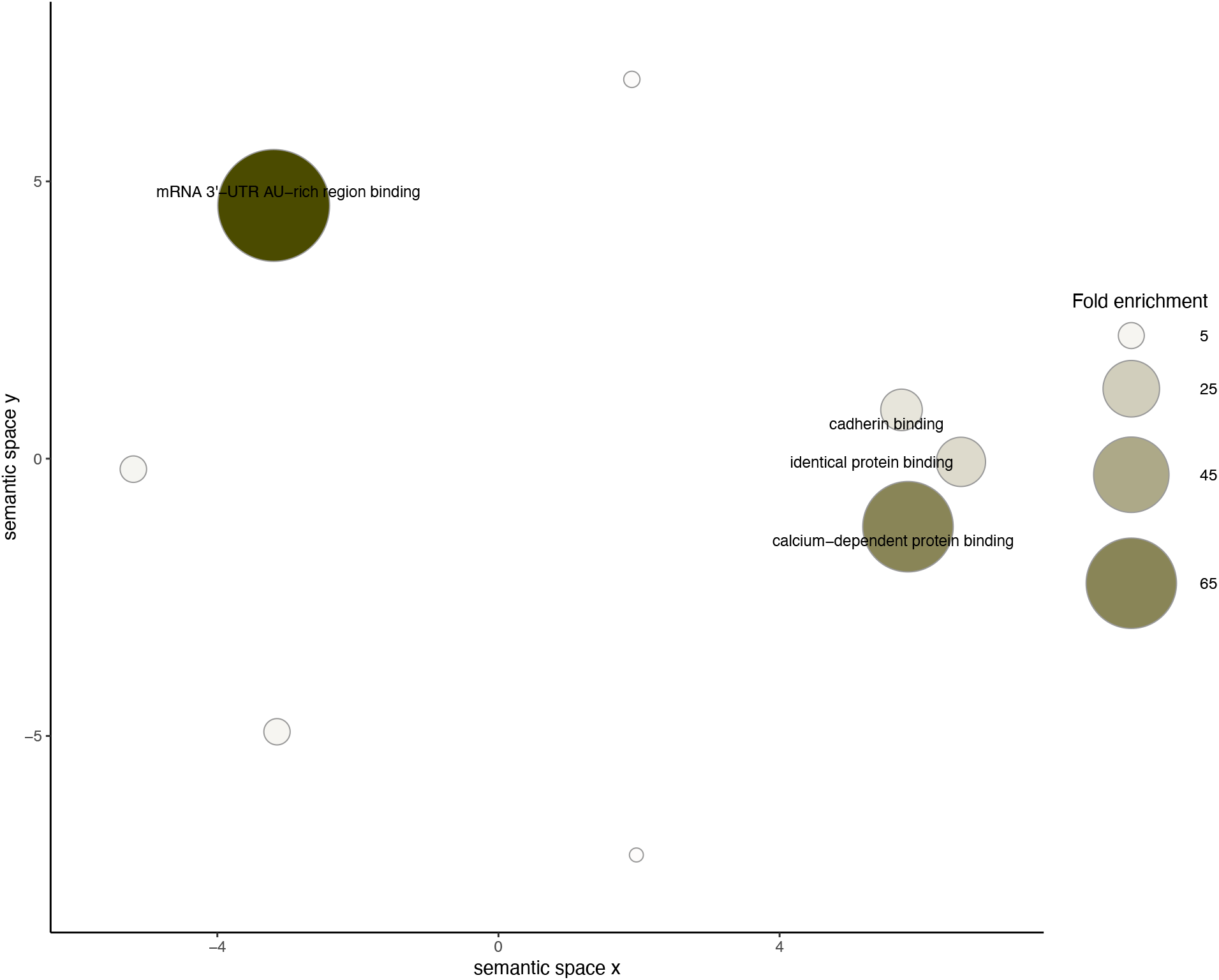
Clustering of statistically significant overrepresented Gene Ontology annotations for molecular functions associated to differences between high and low sociability in guppies. Point size and color provide information on fold enrichment value from the statistical overrepresentation test performed in PANTHER^1^. Terms with fold enrichment lower than eight are represented but not described in text. Axes have no intrinsic meaning and are based on multidimensional scaling which cluster terms based on semantic similarities^2^.

**Figure S4.**
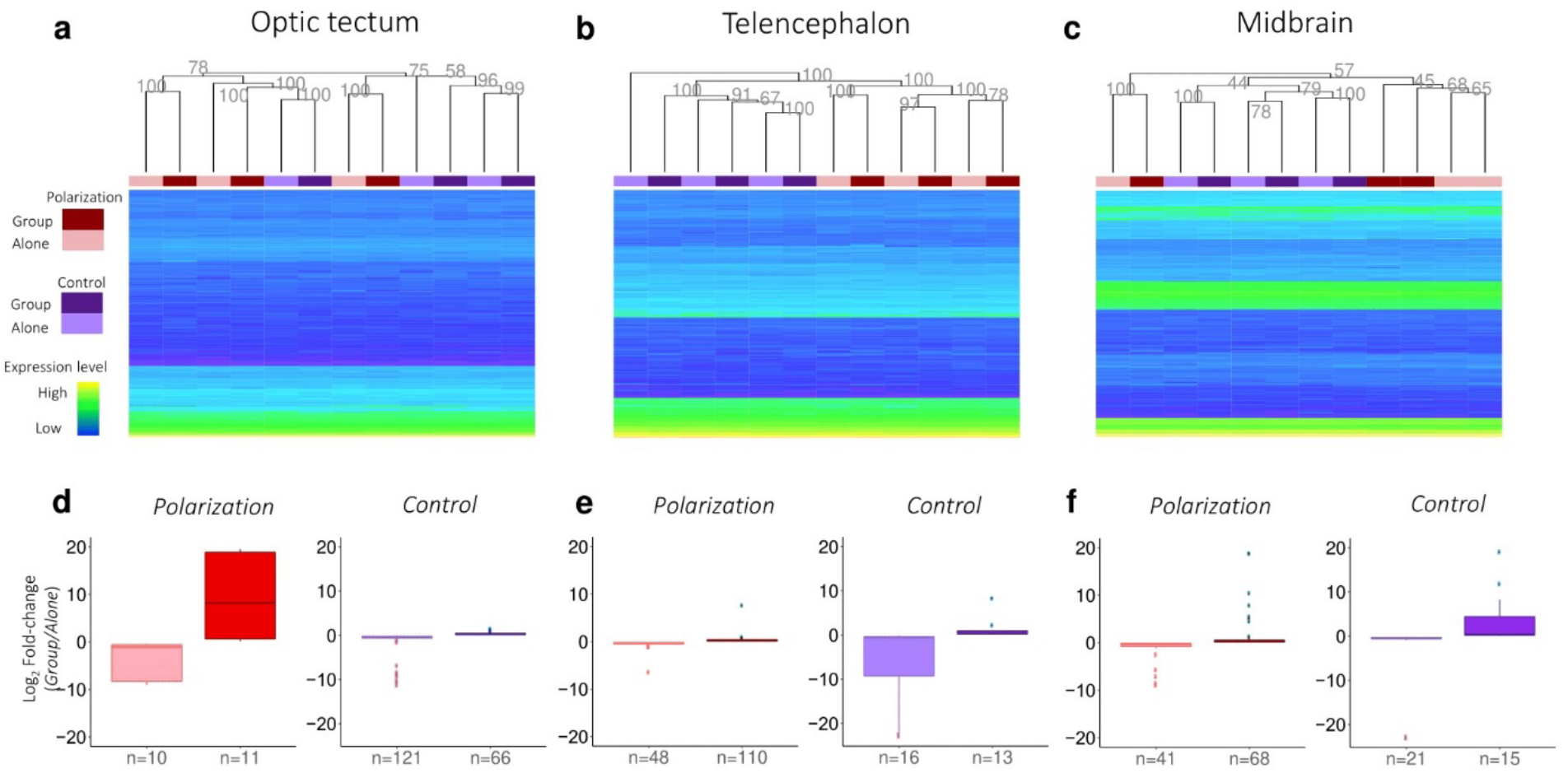
**Top panels:** Hierarchical clustering of gene expression across all significantly expressed genes in the optic tectum (a), telencephalon (b) and midbrain (c) for polarization-selected selection and control lines. Values on top of nodes correspond to Approximately Unbiased bootstrap values. Heatmap represents expression of 500 randomly selected genes. **Bottom panels:** Boxplots of average log2 fold-change for all genes differentially expressed genes between the *Alone* and the *Group* conditions in polarization-selected and control lines for optic tectum (d), telencephalon (e) and midbrain (f). Boxes correspond to 25^th^ - 75th percentiles. Sample sizes on the x-axis indicate number of genes with relative up- or down-regulation.

**Figure S5.**
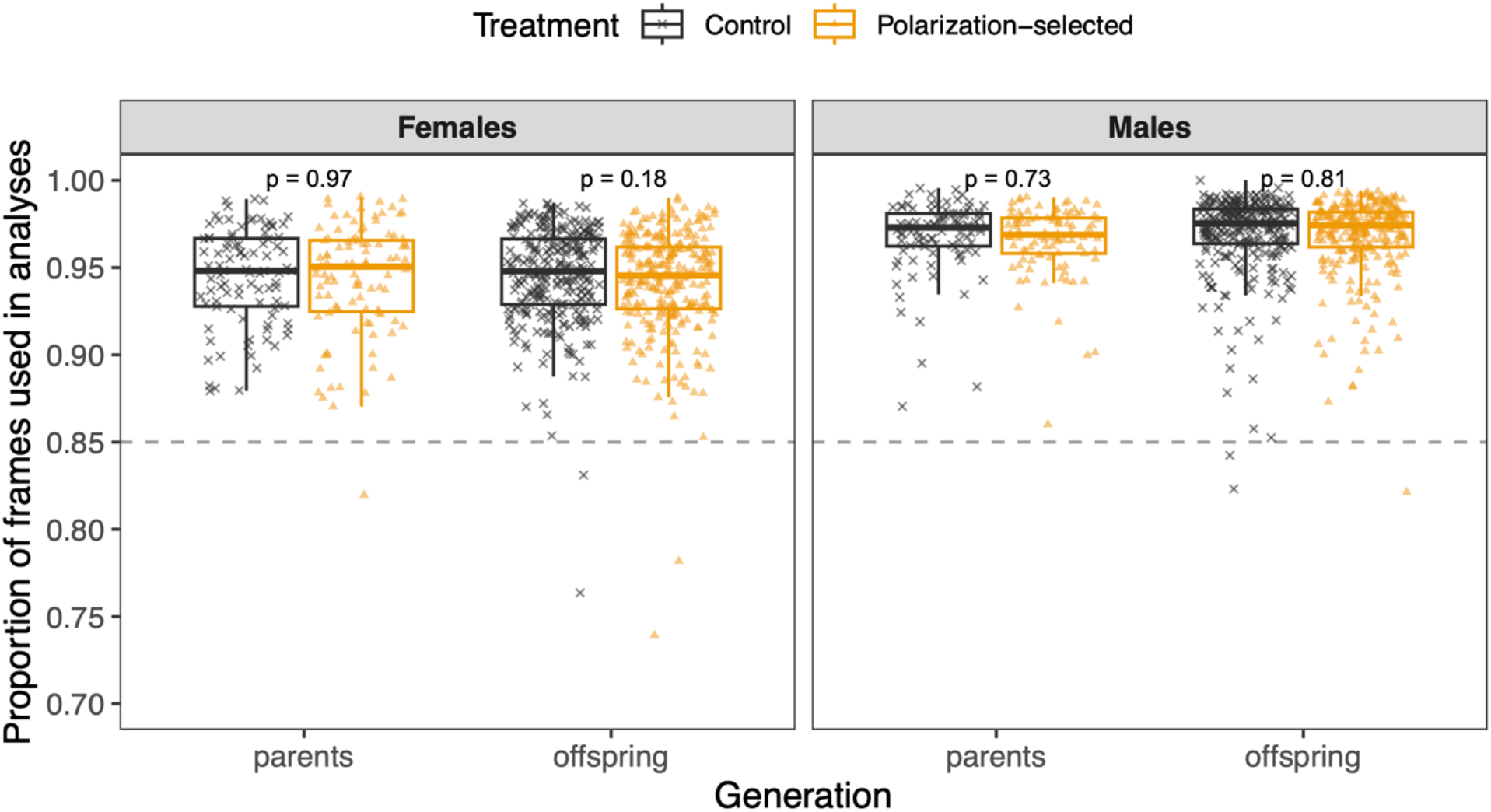
Proportion of frames obtained from idTracker software in tracking from videos recorded for groups of seven wild-type guppies and one individual from either polarization-selected or control selection lines. Tracking data was used next for the quantification of alignment, attraction and speed in parents and six offspring of 195 families. We found no significant differences for the proportion of frames used for videos used for polarization-selected and control lines in a Linear Mixed Model that included generation, sex and selection line as fixed factors (LMM_frames_: line: F= 0.82; df = 1464; p = 0.37). Horizontal lines indicate medians, boxes indicate the interquartile range, and whiskers indicate all points within 1.5 times the interquartile range. P-values in top position of each comparison indicate values for statistical contrasts of the model by sex and generation. Consistency in identity assigned by idTracker was verified in trials with less than 85% frames tracked (dashed line).

## Supplementary Tables for

**Table S1.** Statistical results for a Linear Mixed Model comparing the alignment and attraction of polarization-selected and control female guppies when swimming with groups of same-sex non-kin conspecifics.

| **Alignment to group** |  |  |  |  |  |
| --- | --- | --- | --- | --- | --- |
| *Predictors* | *Estimates* | *CI* | *Statistic* | *p* | *df* |
| (Intercept) | 0.19 | 0.13 – 0.24 | 6.78 | **<0.001** | 1385.81 |
| Selection line [polarization] | 0.01 | 0.00 – 0.01 | 2.27 | **0.047** | 9.68 |
| Sex [males] | -0.01 | -0.01 – -0.00 | -3.02 | **0.003** | 690.08 |
| Generation [parental] | -0.02 | -0.02 – -0.01 | -10.13 | **<0.001** | 1141.24 |
| Average alignment of group | 0.79 | 0.72 – 0.86 | 22.71 | **<0.001** | 1436.72 |
| Average speed of group | 0.00 | 0.00 – 0.00 | 0.20 | 0.842 | 1429.55 |
| Selection line [polarization] *<br>Sex [males] | 0.00 | -0.00 – 0.00 | 0.20 | 0.842 | 1429.55 |
| ICC | 0.12 |  |  |  |  |
| N<sub>line</sub> | 2 |  |  |  |  |
| N<sub>rep</sub> | 3 |  |  |  |  |
| N<sub>testnr</sub> | 8 |  |  |  |  |
| N<sub>sex</sub> | 2 |  |  |  |  |
| N<sub>family</sub> | 195 |  |  |  |  |
| Observations | 1486 |  |  |  |  |
| Marginal R<sup>2</sup> / Conditional R<sup>2</sup> | 0.657 / 0.697 |  |  |  |  |

| **Nearest neighbour distance (attraction)** |  |  |  |  |  |
| --- | --- | --- | --- | --- | --- |
| *Predictors* | *Estimates* | *CI* | *Statistic* | *p* | *df* |
| (Intercept) | 2.64 | 2.56 – 2.71 | 65.62 | **<0.001** | 565.12 |
| Selection line [polarization] | -0.04 | -0.08 – -0.00 | -2.33 | **0.044** | 9.34 |
| Sex [males] | 0.01 | -0.02 – 0.04 | 0.55 | 0.583 | 448.53 |
| Generation [parental] | 0.12 | 0.10 – 0.14 | 11.31 | **<0.001** | 984.50 |
| Average alignment of group | 0.02 | 0.02 – 0.02 | 27.31 | **<0.001** | 1408.27 |
| Average speed of group | 0.00 | 0.00 – 0.00 | 3.74 | **<0.001** | 566.82 |
| Selection line [polarization] * Sex<br>[males] | 0.03 | -0.02 – 0.07 | 1.24 | 0.217 | 371.59 |
| ICC | 0.13 |  |  |  |  |
| N<sub>line</sub> | 2 |  |  |  |  |
| N<sub>rep</sub> | 3 |  |  |  |  |
| N<sub>testnr</sub> | 8 |  |  |  |  |
| N<sub>sex</sub> | 2 |  |  |  |  |
| N<sub>family</sub> | 195 |  |  |  |  |
| Observations | 1485 |  |  |  |  |
| Marginal R<sup>2</sup> / Conditional R<sup>2</sup> | 0.471 / 0.538 |  |  |  |  |

**Table S2.** Independent contrasts for comparisons between polarization-selected (P) and control (C) guppies in their alignment and nearest neighbor distance (attraction) when swimming with groups of same-sex non-kin conspecifics.

Alignment
| *contrast* | *sex* | *estimate* | *SE* | *df* | *t.ratio* | *p.value* |
| --- | --- | --- | --- | --- | --- | --- |
| C - P | females | -0.006 | 0.002 | 9.68 | -2.269 | **0.047** |
| C - P | males | -0.003 | 0.002 | 9.56 | 1.378 | 0.20 |

Attraction
| *contrast* | *treatment* | *estimate* | *SE* | *df* | *t.ratio* | *p.value* |
| --- | --- | --- | --- | --- | --- | --- |
| C - P | females | 0.042 | 0.001 | 9.34 | 2.331 | **0.043** |
| C - P | males | 0.015 | 0.002 | 9.26 | 0.883 | 0.40 |

**Table S3.**
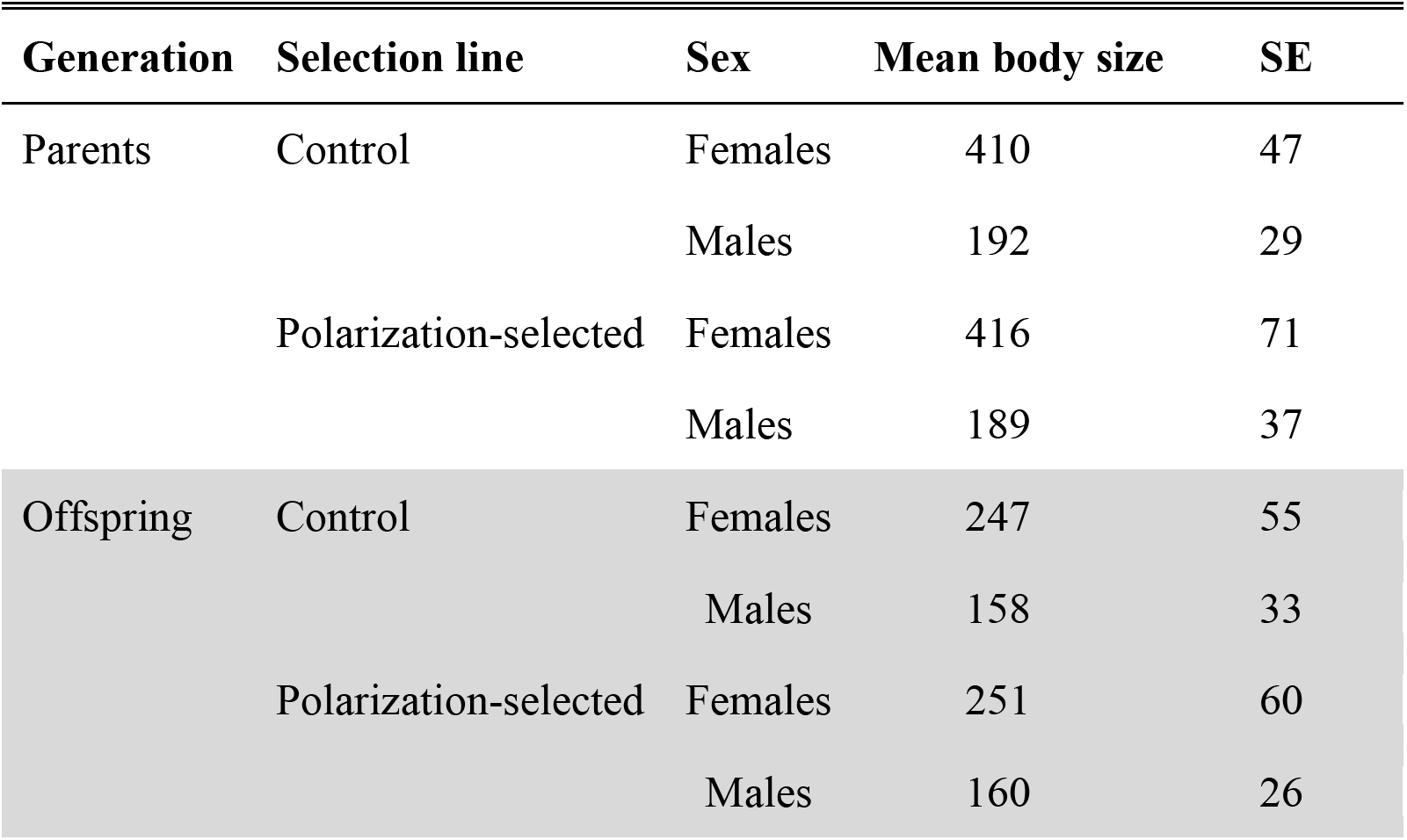
Mean body size of guppies used to estimate the heritability of alignment and attraction at the time of testing across sexes and generations. Values are provided in number of pixels obtained from idTracker data using a custom script implemented in Matlab (v2020a) that extracts body size data accounting for changes in apparent size between the middle and edge of the experimental arena used for video recordings.

**Table S4.** Results from animal models only accounting for same-sex pedigree relationships. We obtained heritability estimates by taking the ratio of the additive genetic variance to the total phenotypic variance (additive genetic variance + group variance + population variance + residual variance) in each independent model.

ALIGNMENT
| FEMALES |  |  |  |  |  |
| --- | --- | --- | --- | --- | --- |
| Parameter group | Parameter | Estimate | Est.Error | Q2.5 | Q97.5 |
| Fixed effects | Intercept | 0.504 | 0.074 | 0.359 | 0.648 |
| Fixed effects | linesocial | 0.157 | 0.102 | -0.04 | 0.355 |
| Fixed effects | generationparents | -0.277 | 0.058 | -0.39 | -0.163 |
| Fixed effects | speed_std | 0.396 | 0.023 | 0.35 | 0.441 |
| Additive genetic variance | Intercept | 0.162 | 0.042 | 0.086 | 0.25 |
| Group | Intercept | 0.005 | 0.006 | 0 | 0.02 |
| Population | Intercept | 0.01 | 0.027 | 0 | 0.064 |
| Residual variance | NA | 0.304 | 0.037 | 0.234 | 0.379 |
| Heritability |  | 0.336 | 0.079 | 0.183 | 0.494 |

| MALES |  |  |  |  |  |
| --- | --- | --- | --- | --- | --- |
| Parameter group | Parameter | Estimate | Est.Error | Q2.5 | Q97.5 |
| Fixed effects | Intercept | -0.452 | 0.125 | -0.694 | -0.213 |
| Fixed effects | linesocial | 0.093 | 0.172 | -0.231 | 0.425 |
| Fixed effects | generationparents | -0.502 | 0.071 | -0.641 | -0.362 |
| Fixed effects | speed_std | 0.298 | 0.039 | 0.221 | 0.376 |
| Additive genetic variance | Intercept | 0.031 | 0.026 | 0.000 | 0.091 |
| Group | Intercept | 0.04 | 0.016 | 0.013 | 0.074 |
| Population | Intercept | 0.039 | 0.091 | 0.001 | 0.207 |
| Residual variance | NA | 0.390 | 0.033 | 0.323 | 0.453 |
| Heritability |  | 0.063 | 0.052 | 0.000 | 0.184 |

ATTRACTION
| FEMALES |  |  |  |  |  |
| --- | --- | --- | --- | --- | --- |
| Parameter group | Parameter | Estimate | Est.Error | Q2.5 | Q97.5 |
| Fixed effects | Intercept | -0.614 | 0.099 | -0.811 | -0.424 |
| Fixed effects | linesocial | -0.088 | 0.138 | -0.35 | 0.179 |
| Fixed effects | generationparents | 0.251 | 0.070 | 0.113 | 0.390 |
| Fixed effects | speed_std | -0.109 | 0.024 | -0.157 | -0.062 |
| Additive genetic variance | Intercept | 0.094 | 0.038 | 0.025 | 0.174 |
| Group | Intercept | 0.038 | 0.015 | 0.011 | 0.071 |
| Population | Intercept | 0.023 | 0.061 | 0.000 | 0.133 |
| Residual variance | NA | 0.355 | 0.039 | 0.280 | 0.435 |
| Heritability |  | 0.185 | 0.072 | 0.049 | 0.336 |

MALES
| Parameter group | Parameter | Estimate | Est.Error | Q2.5 | Q97.5 |
| --- | --- | --- | --- | --- | --- |
| Fixed effects | Intercept | 0.476 | 0.134 | 0.212 | 0.746 |
| Fixed effects | linesocial | -0.025 | 0.181 | -0.389 | 0.345 |
| Fixed effects | generationparents | 0.614 | 0.088 | 0.440 | 0.785 |
| Fixed effects | speed_std | 0.054 | 0.046 | -0.037 | 0.145 |
| Additive genetic variance | Intercept | 0.137 | 0.049 | 0.047 | 0.242 |
| Group | Intercept | 0.075 | 0.023 | 0.036 | 0.126 |
| Population | Intercept | 0.046 | 0.097 | 0.000 | 0.250 |
| Residual variance | NA | 0.454 | 0.050 | 0.359 | 0.556 |
| **Heritability** |  | 0.194 | 0.069 | 0.065 | 0.337 |

**Table S5.** Results from animal models with full pedigree relationships. We obtained heritability estimates for each sex by taking the ratio of the additive genetic variance of each sex to the total phenotypic variance (additive genetic variance + group variance + population variance + residual variance) in each independent model.

ALIGNMENT
| Parameter group | Parameter | Estimate | Est.Error | Q2.5 | Q97.5 |
| --- | --- | --- | --- | --- | --- |
| Fixed effects | Intercept | 0.52 | 0.102 | 0.331 | 0.718 |
| Fixed effects | sexm | -0.961 | 0.063 | -1.08 | -0.838 |
| Fixed effects | linesocial | 0.163 | 0.128 | -0.097 | 0.419 |
| Fixed effects | speed_std | 0.36 | 0.021 | 0.319 | 0.4 |
| Fixed effects | sexm:linesocial | -0.075 | 0.074 | -0.22 | 0.07 |
| Fixed effects | sexf:generationparents | -0.31 | 0.066 | -0.444 | -0.181 |
| Fixed effects | sexm:generationparents | -0.479 | 0.064 | -0.604 | -0.352 |
| Additive genetic variance | sexf | 0.105 | 0.029 | 0.051 | 0.166 |
| Additive genetic variance | sexm | 0.048 | 0.027 | 0.003 | 0.105 |
| Additive genetic variance | cross (sexf x sexm) | 0.583 | 0.270 | 0.021 | 0.974 |
| Group | Intercept | 0.025 | 0.009 | 0.009 | 0.045 |
| Population | Intercept | 0.023 | 0.051 | 0.001 | 0.13 |
| Residual variance | NA | 0.365 | 0.026 | 0.314 | 0.415 |
|  | ♀ Heritability | 0.202 | 0.052 | 0.102 | 0.308 |
|  | ♂ Heritability | 0.103 | 0.057 | 0.007 | 0.224 |
|  | Δ Heritability (♀ - ♂) | 0.098 | 0.053 | 0.202 | -0.005 |

ATTRACTION
| Parameter group | Parameter | Estimate | Est.Error | Q2.5 | Q97.5 |
| --- | --- | --- | --- | --- | --- |
| Fixed effects | Intercept | -0.627 | 0.115 | -0.847 | -0.391 |
| Fixed effects | sexm | 1.09 | 0.075 | 0.942 | 1.23 |
| Fixed effects | linesocial | -0.095 | 0.157 | -0.425 | 0.219 |
| Fixed effects | speed_std | -0.067 | 0.022 | -0.109 | -0.024 |
| Fixed effects | sexm:linesocial | 0.072 | 0.082 | -0.091 | 0.235 |
| Fixed effects | sexf:generationparents | 0.283 | 0.079 | 0.129 | 0.438 |
| Fixed effects | sexm:generationparents | 0.568 | 0.080 | 0.411 | 0.722 |
| Additive genetic variance | sexf | 0.050 | 0.025 | 0.007 | 0.104 |
| Additive genetic variance | sexm | 0.176 | 0.041 | 0.102 | 0.263 |
| Additive genetic variance | sexm | 0.438 | 0.304 | -0.168 | 0.950 |
| Group | Intercept | 0.059 | 0.013 | 0.035 | 0.087 |
| Population | Intercept | 0.032 | 0.056 | 0.002 | 0.172 |
| Residual variance | NA | 0.406 | 0.028 | 0.351 | 0.460 |
|  | ♀ Heritability | 0.092 | 0.044 | 0.013 | 0.188 |
|  | ♂ Heritability | 0.261 | 0.055 | 0.157 | 0.371 |
|  | Δ Heritability (♀ - ♂) | -0.169 | 0.051 | -0.267 | -0.069 |

**Table S6.**
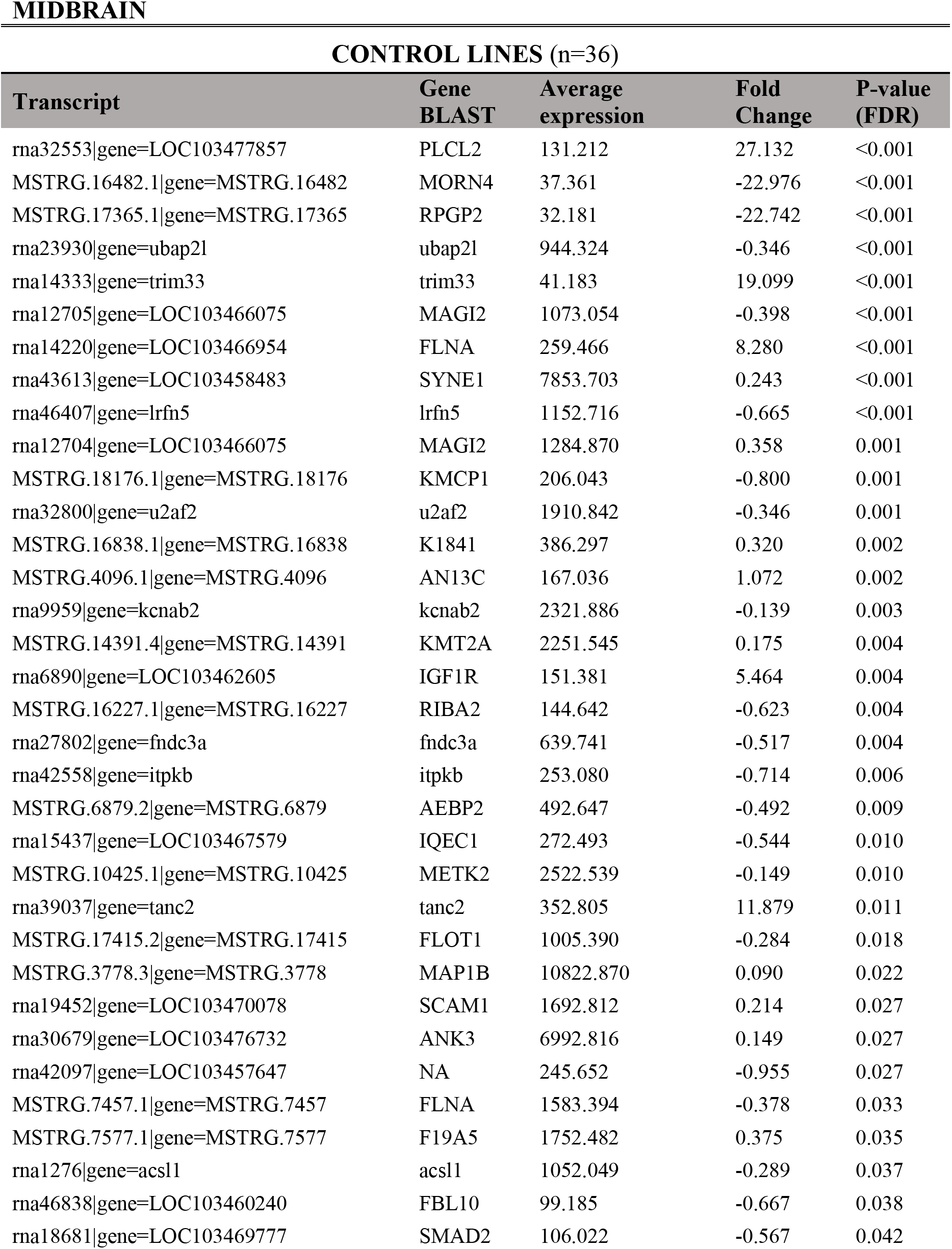
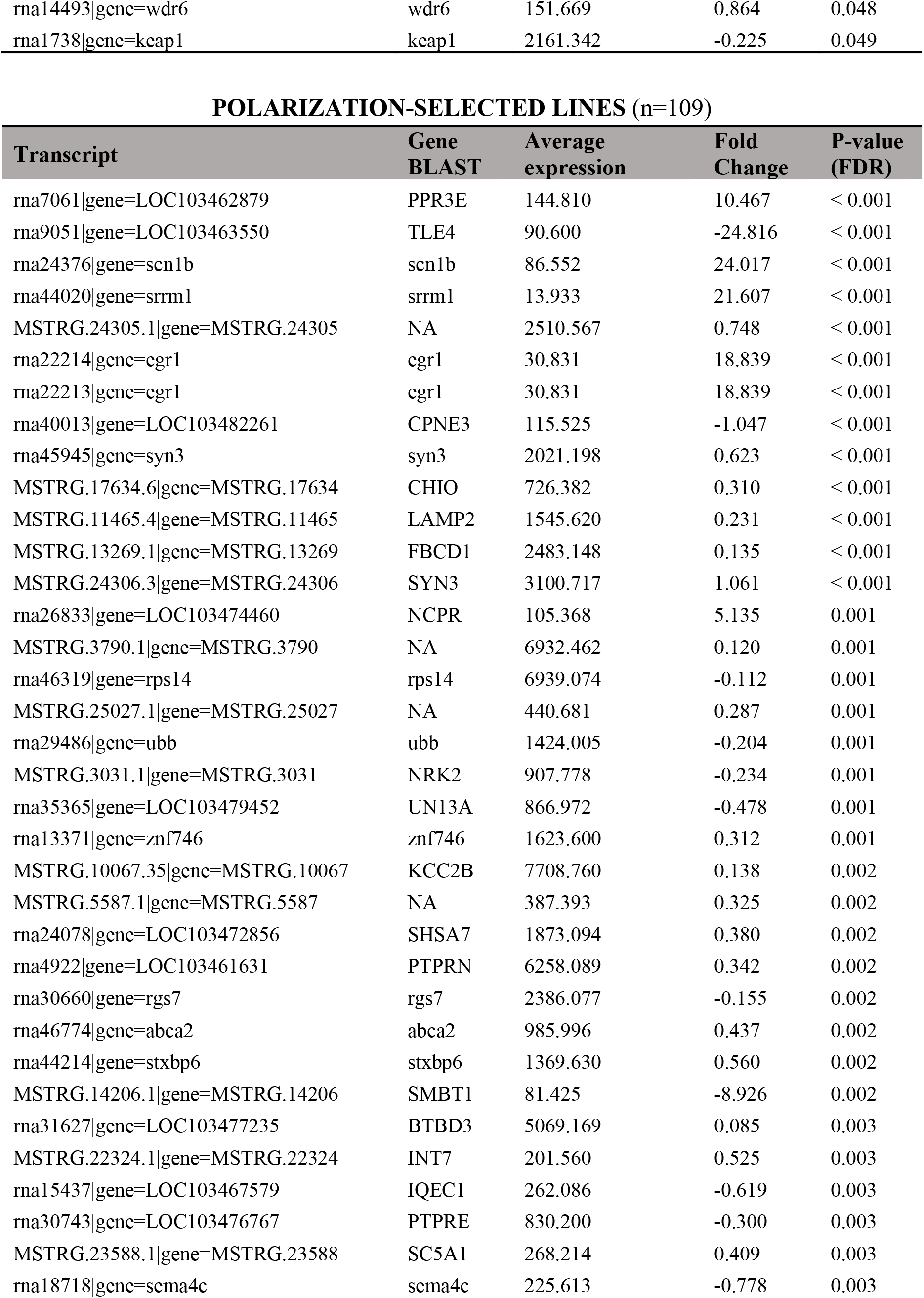
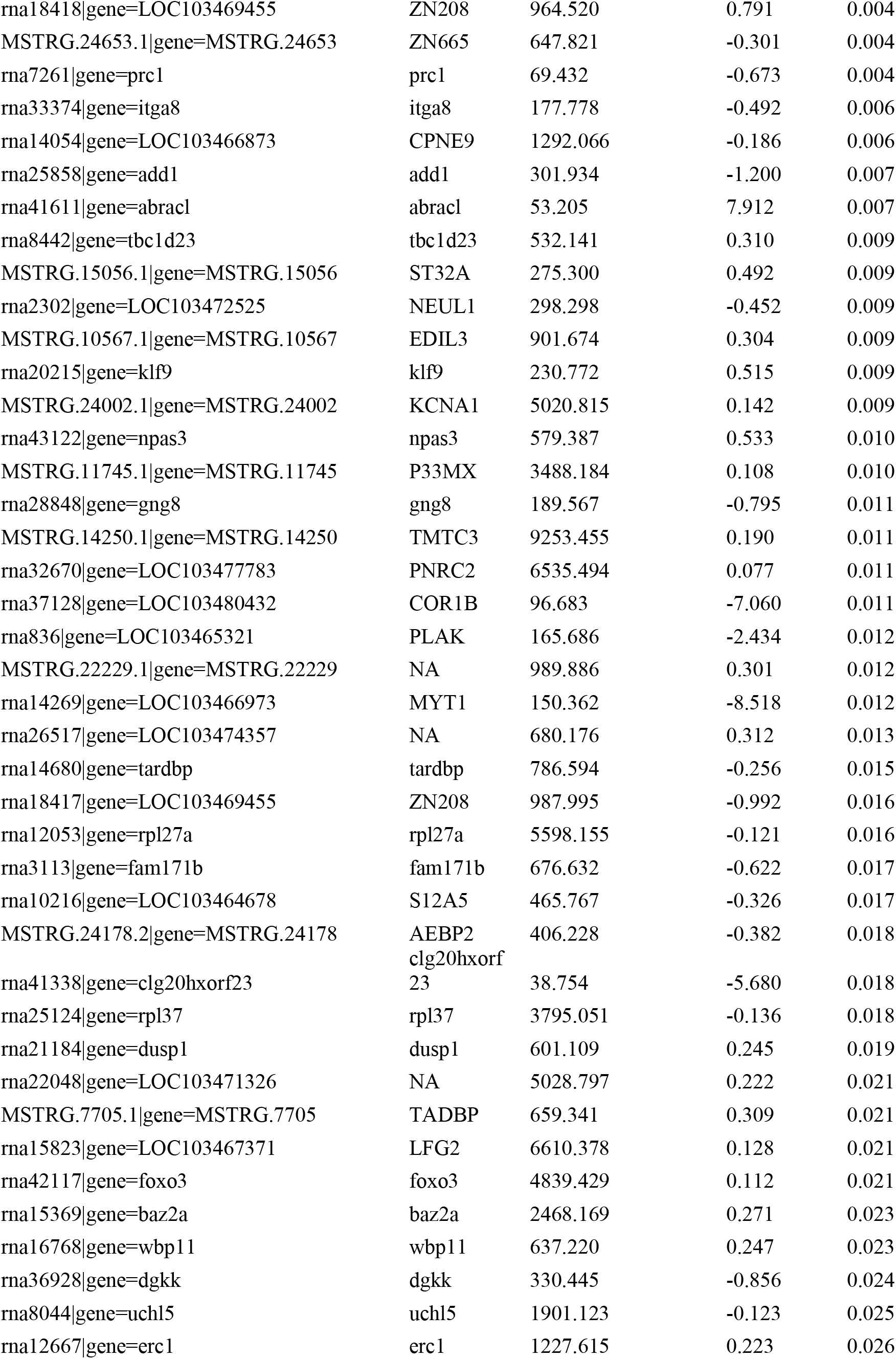
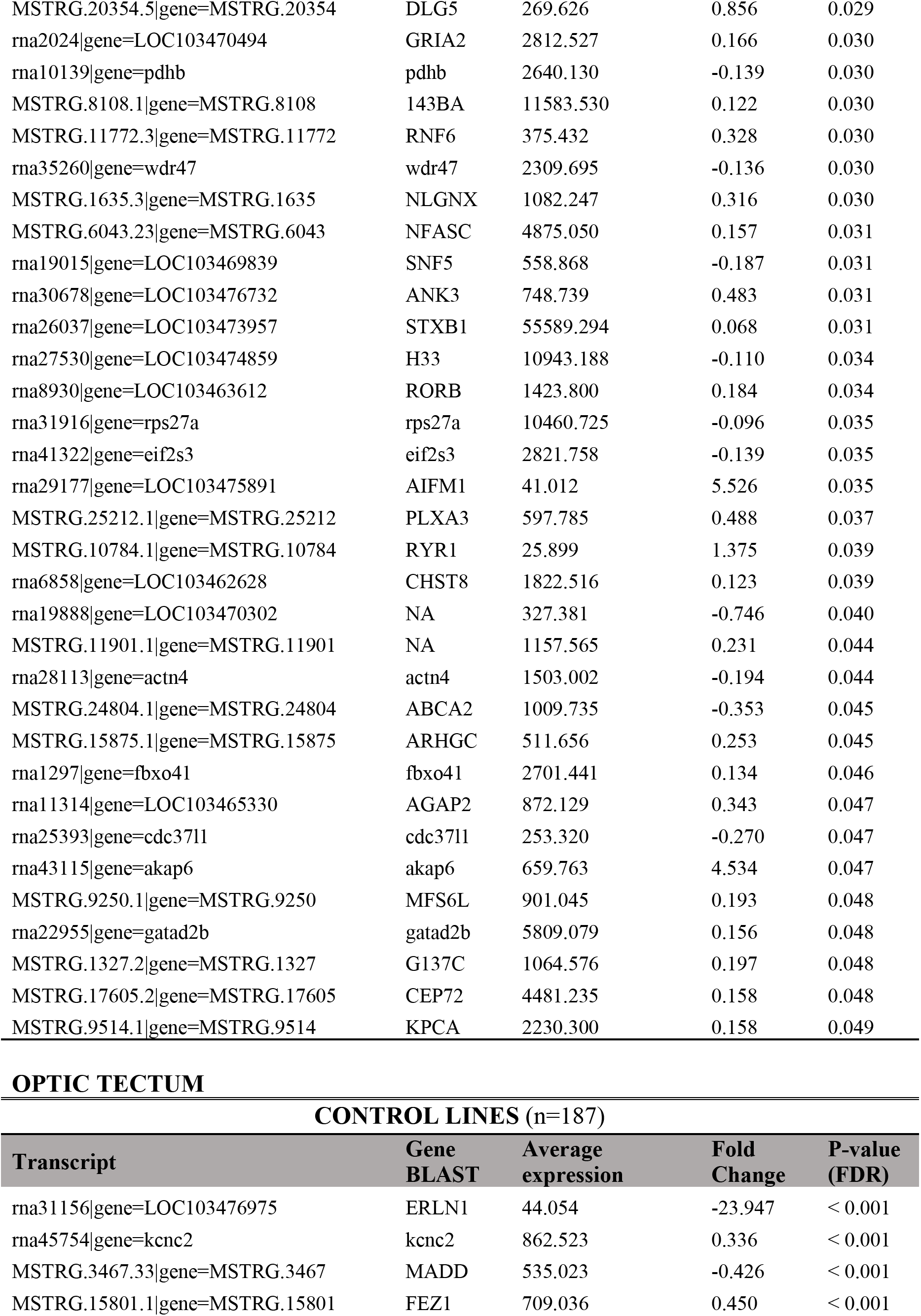
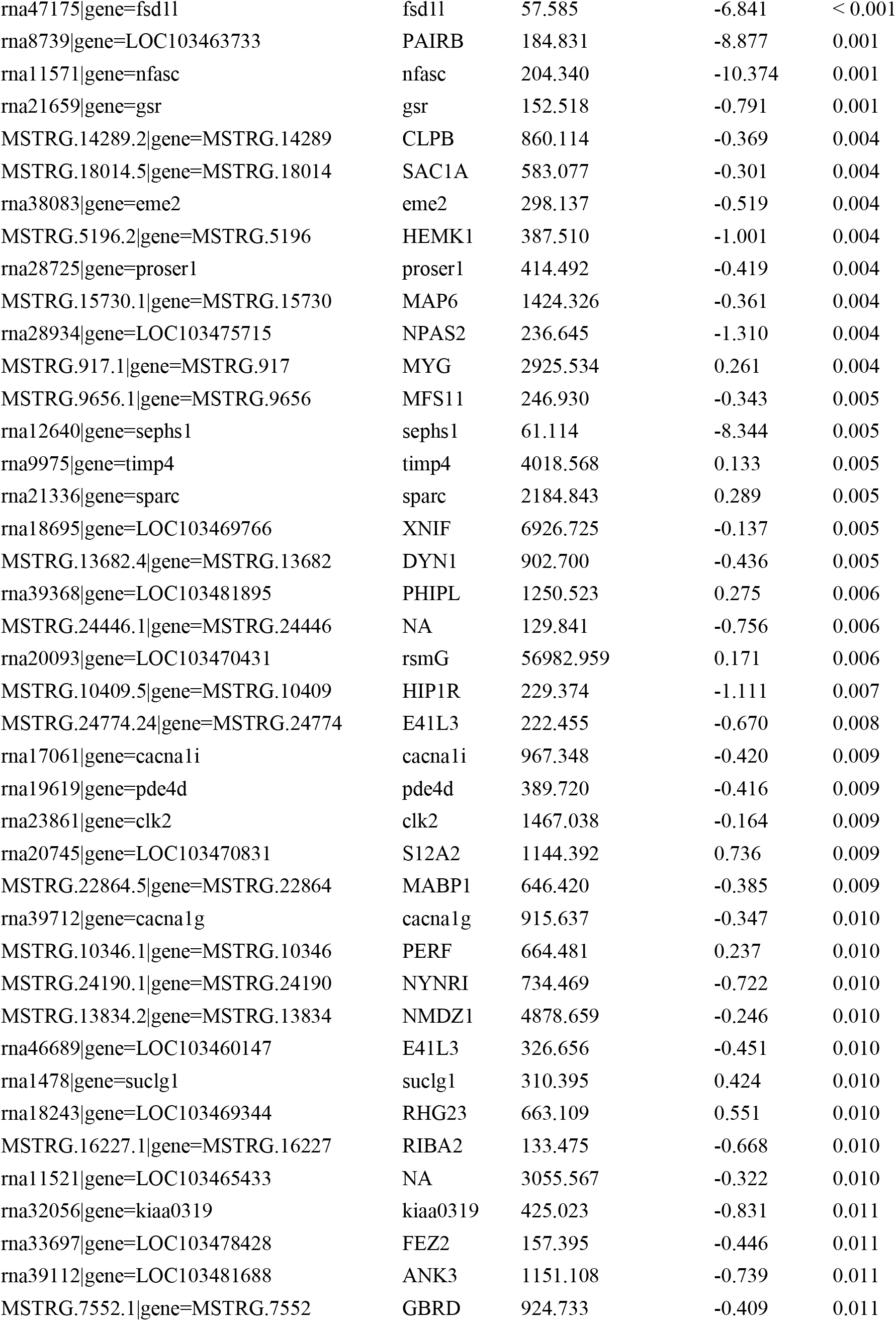
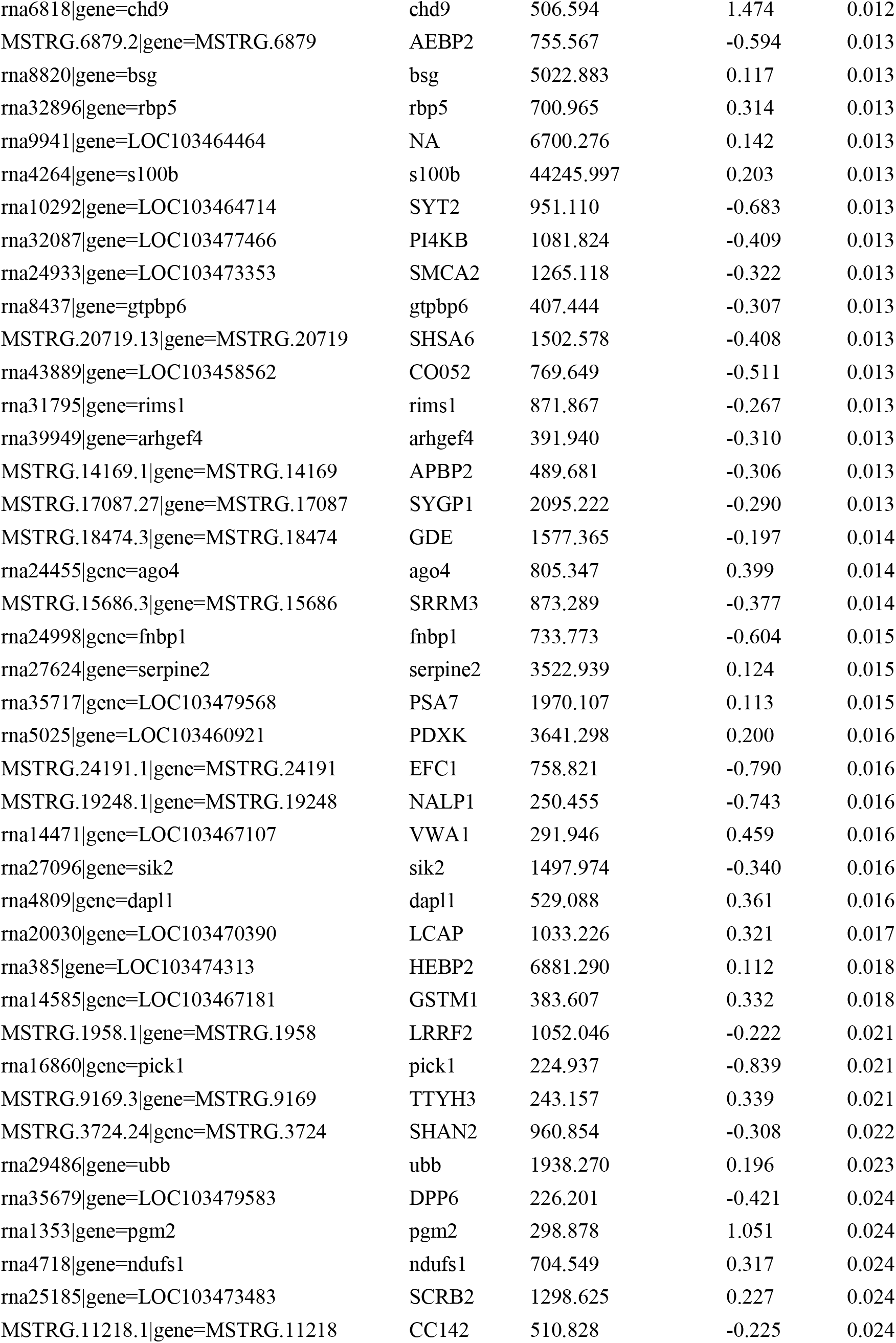
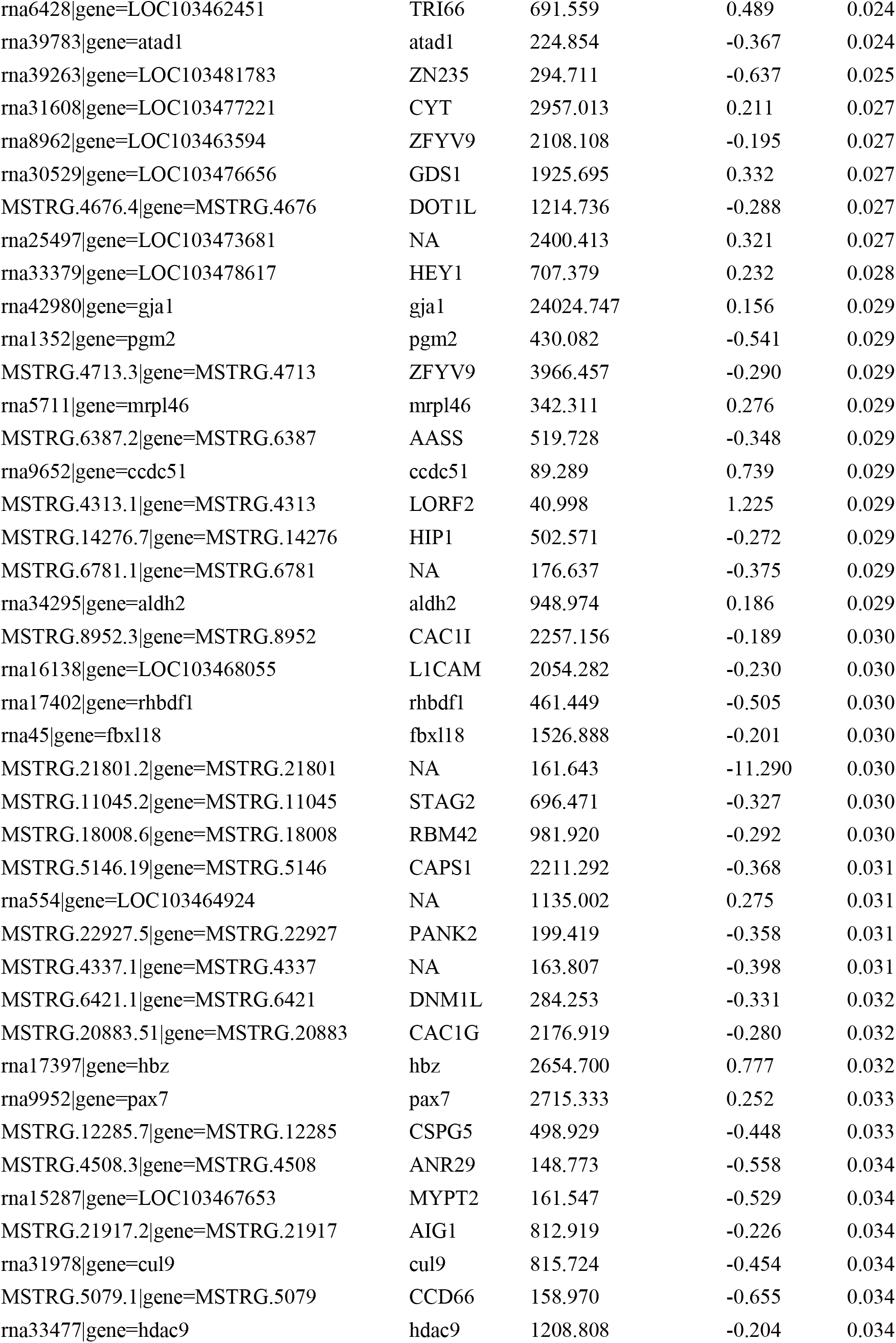
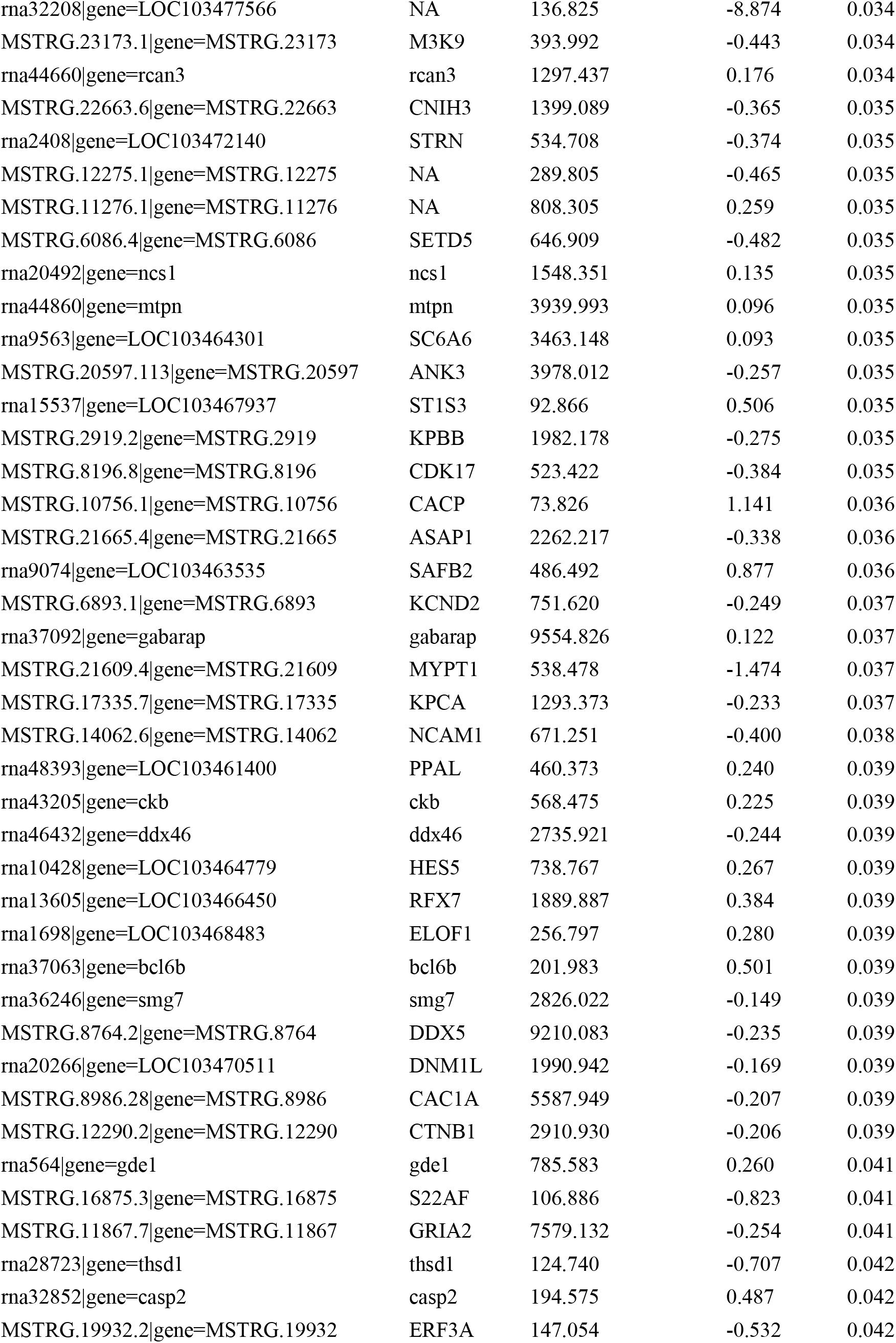
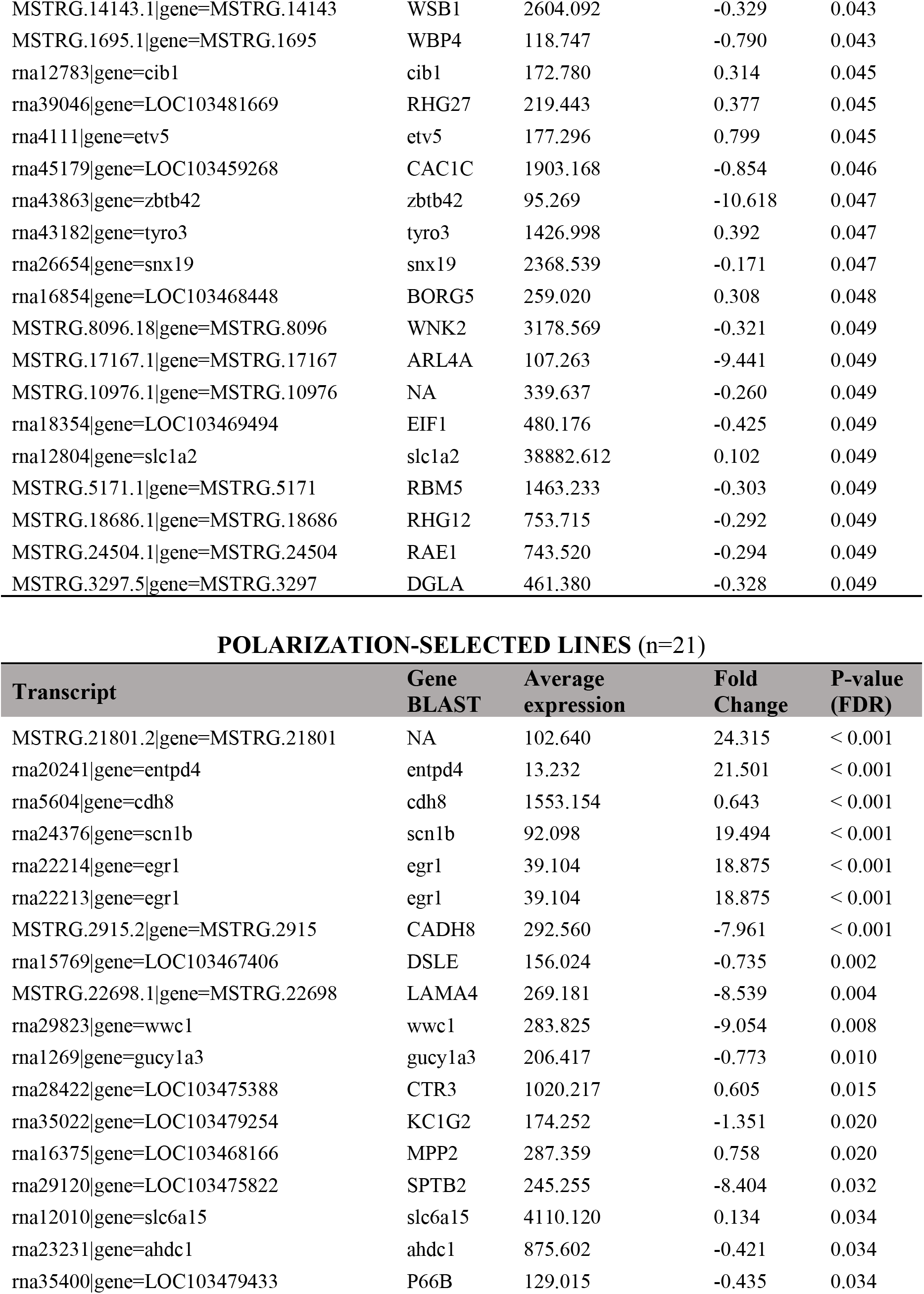
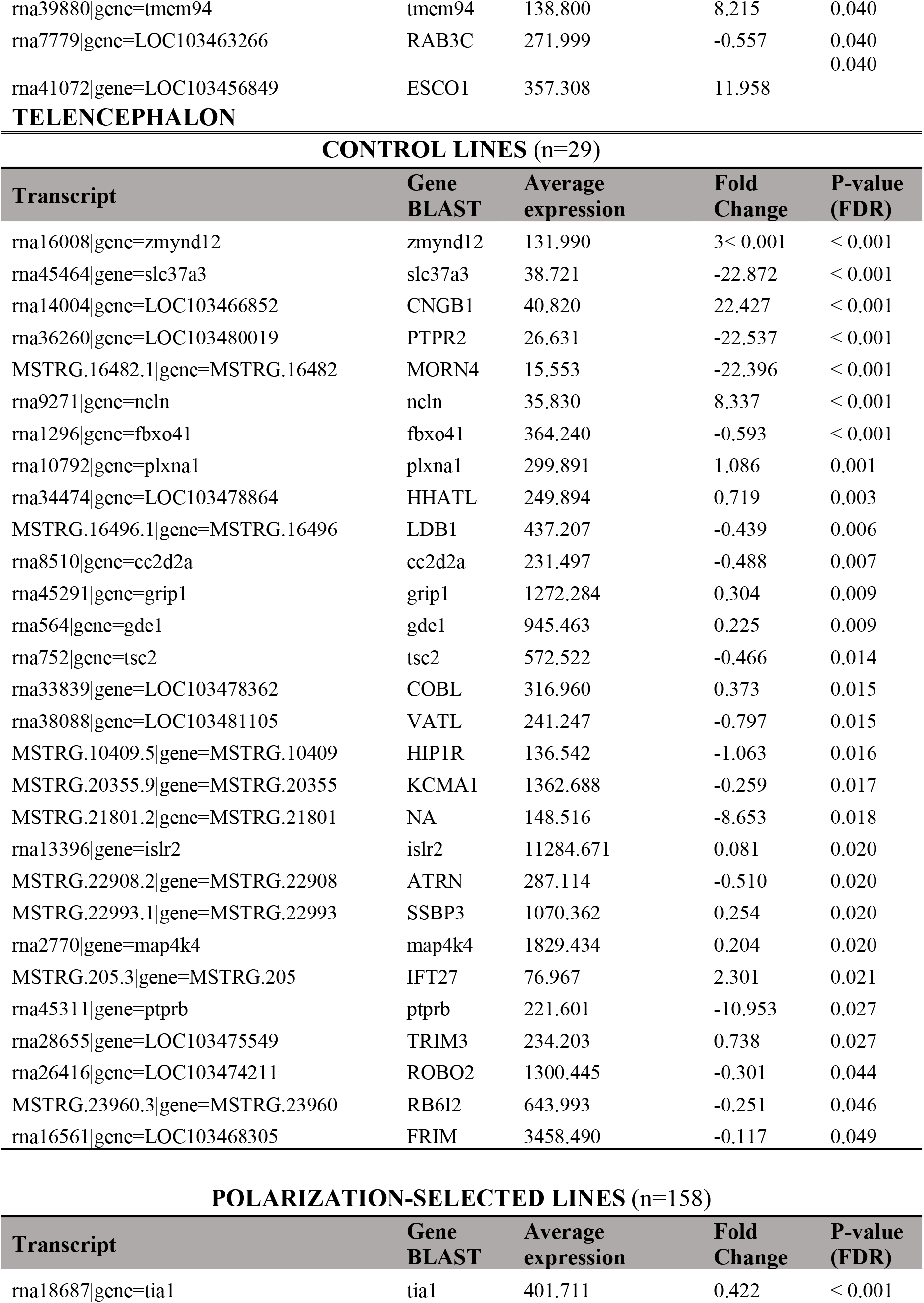
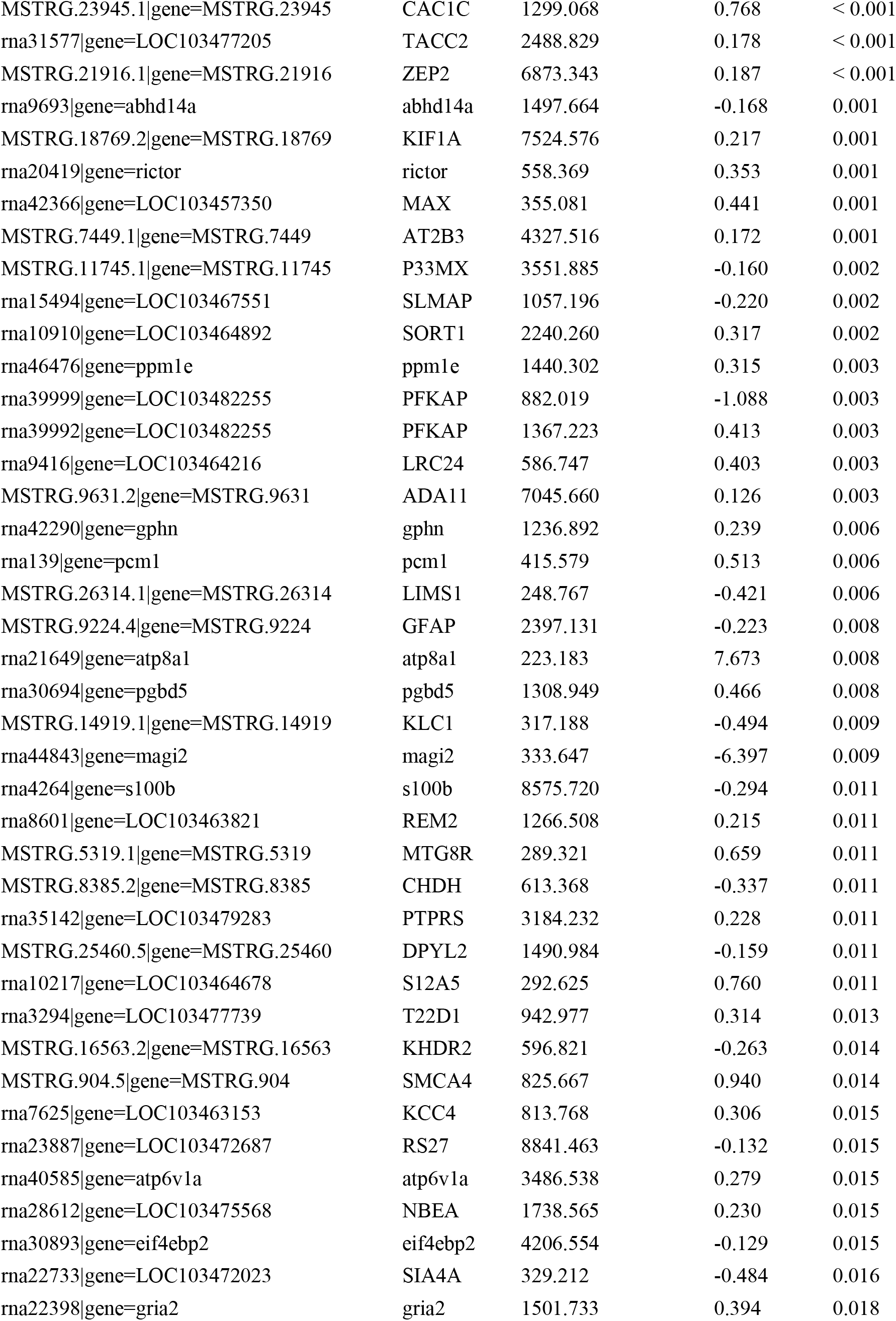
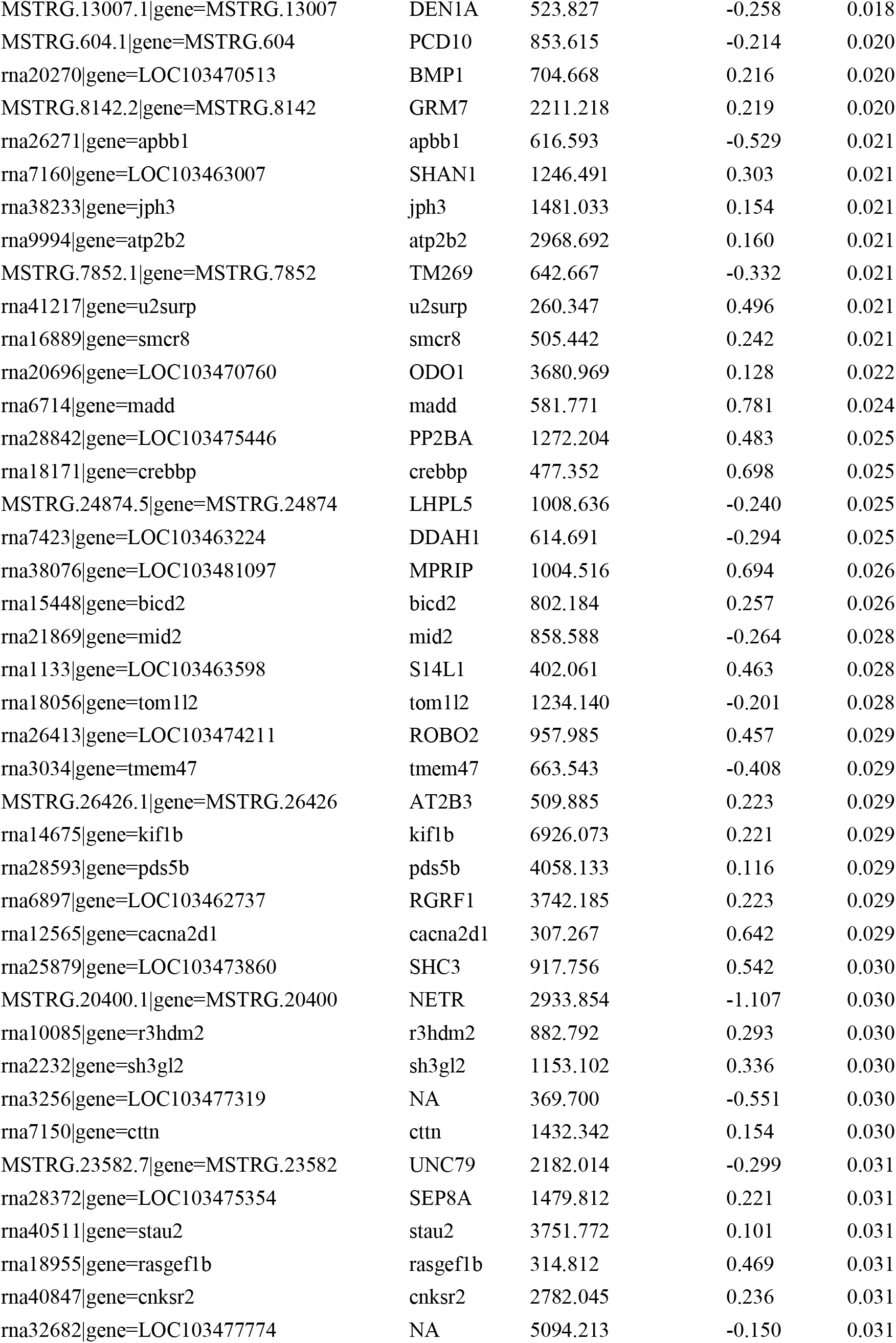
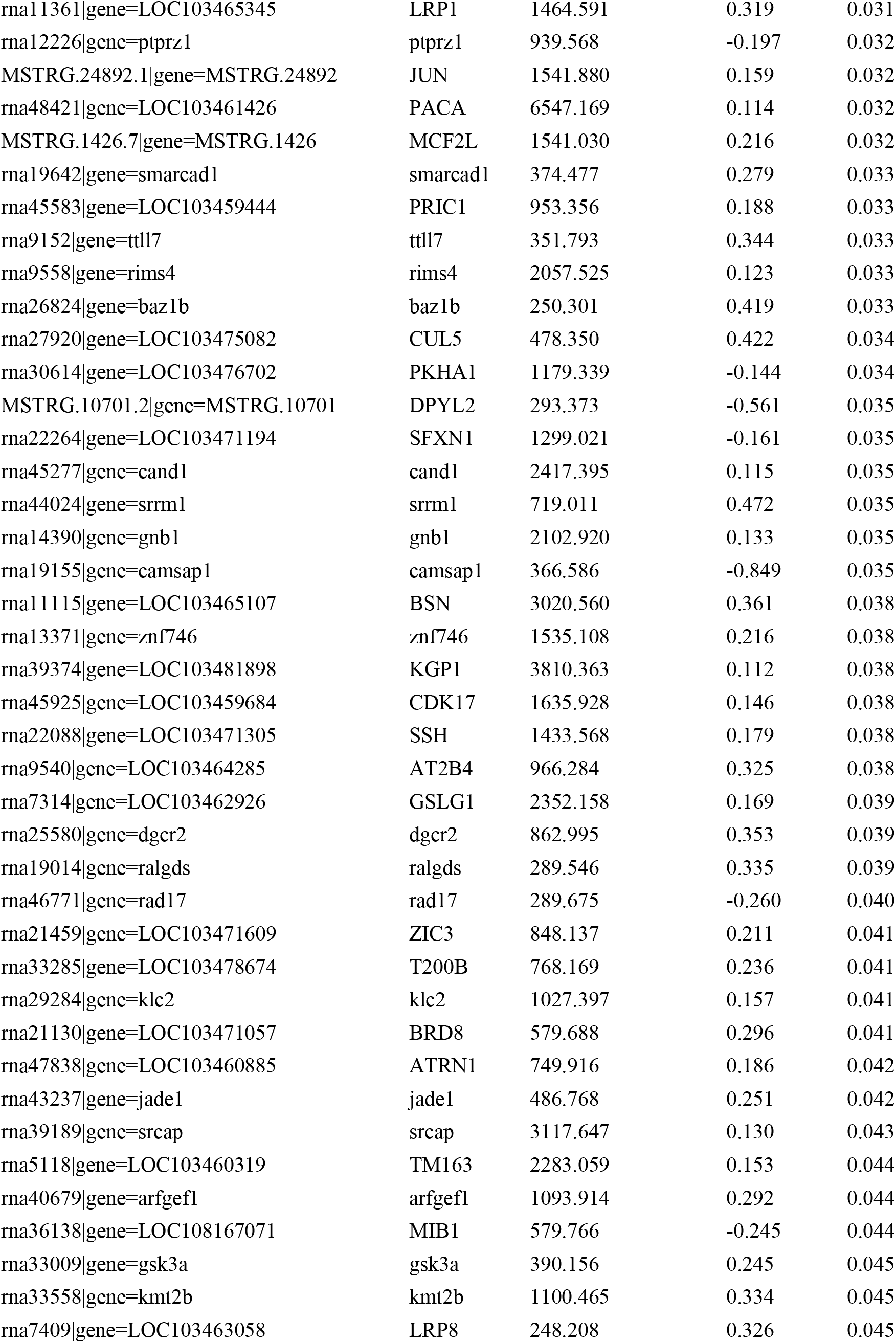
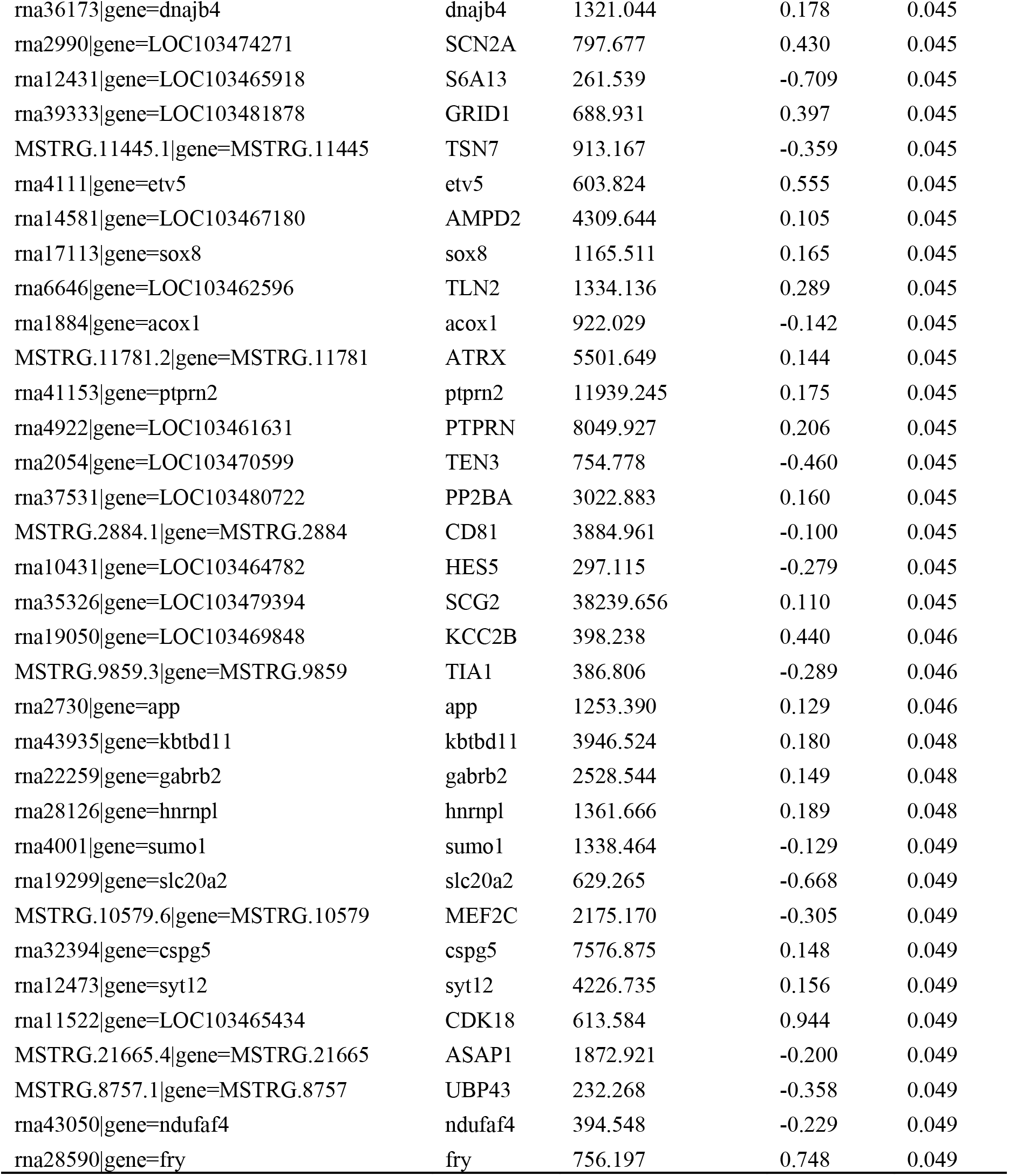
Differentially expressed genes in analyses of the genomic response in midbrain, optic tectum and telencephalon in polarization-selected and control lines of female guppies when swimming with a group of conspecifics versus when swimming alone.

**Table S7.**
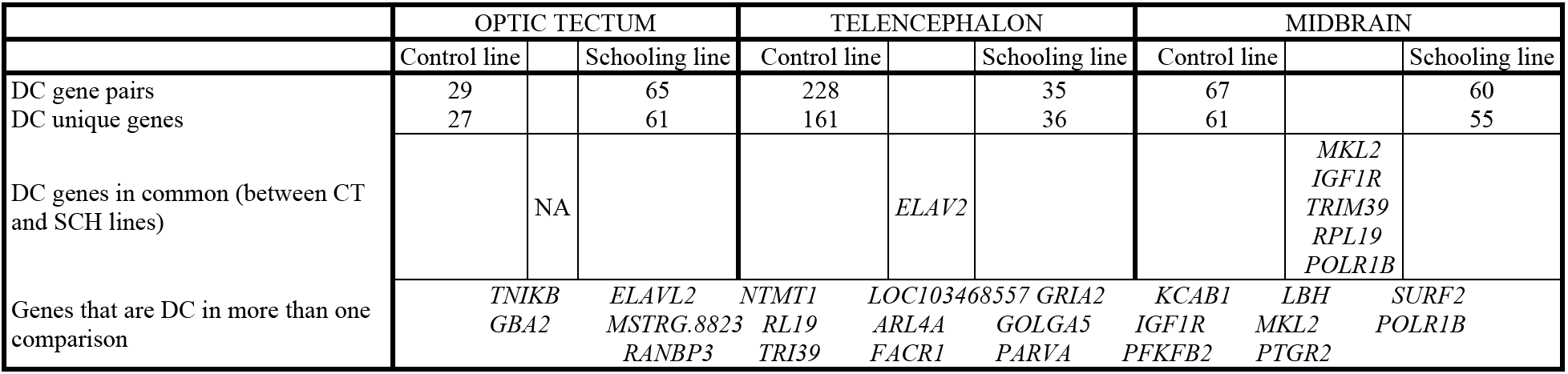
Summary of differentially co-expressed gene pair results

**Table S8.**
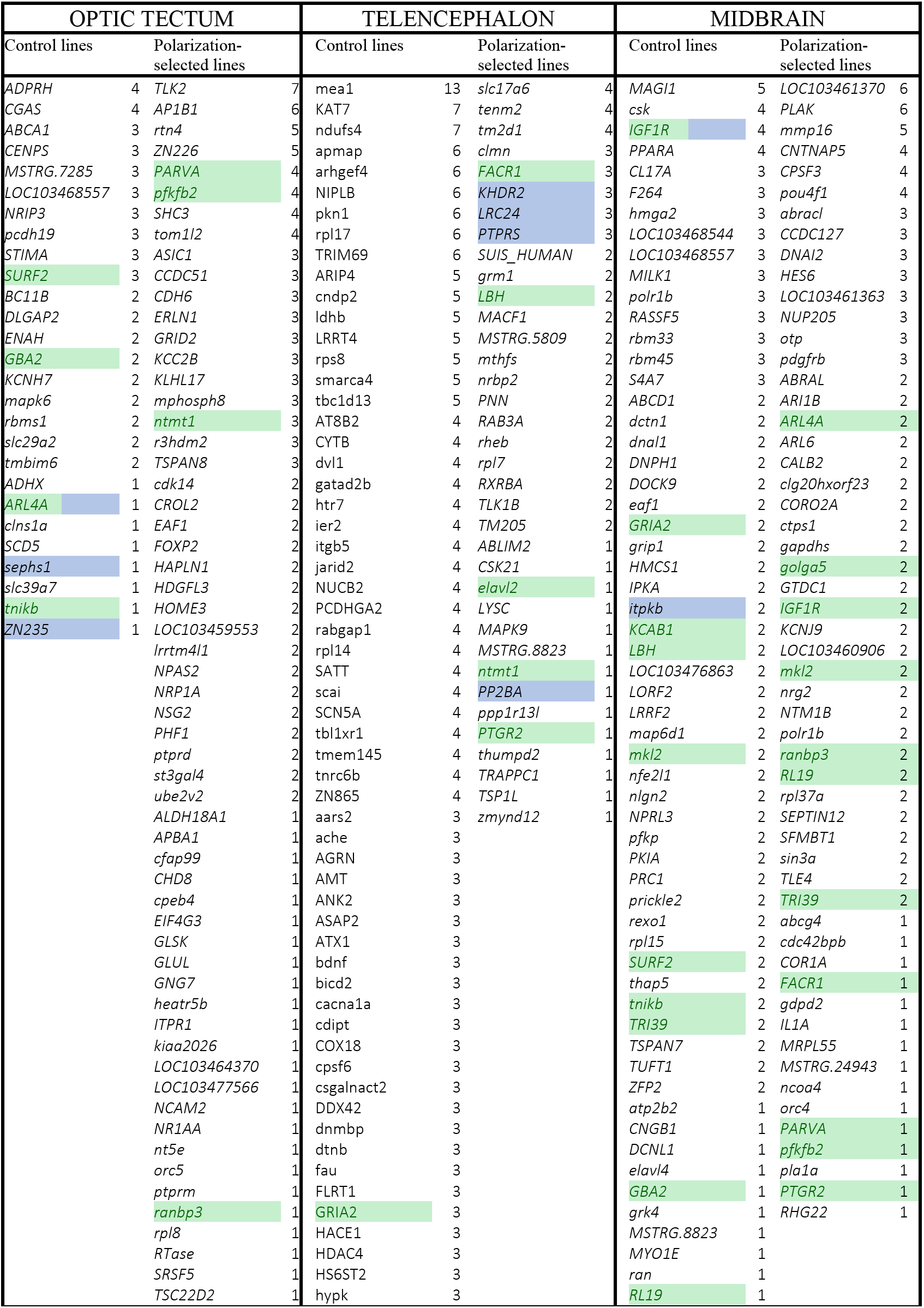
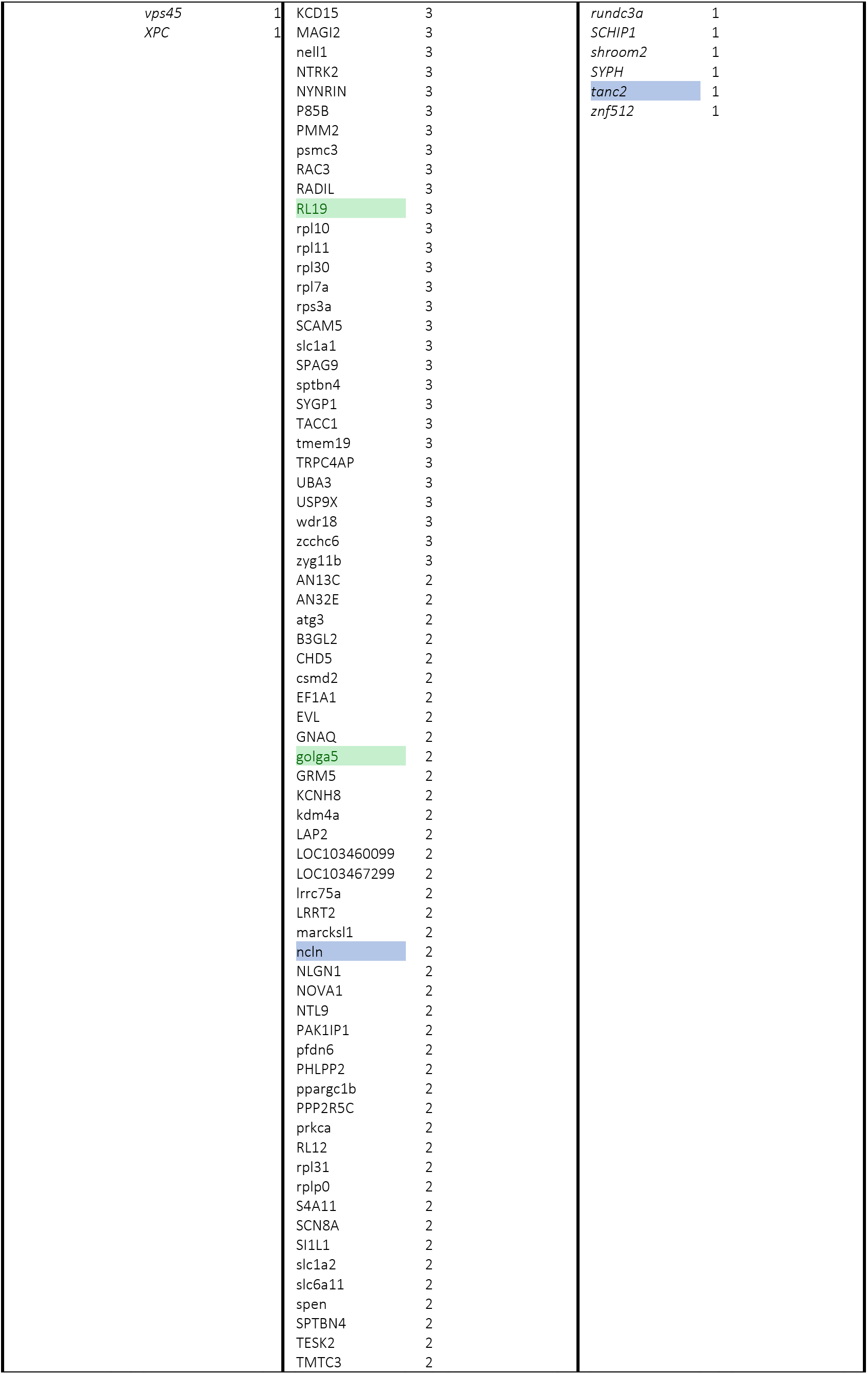
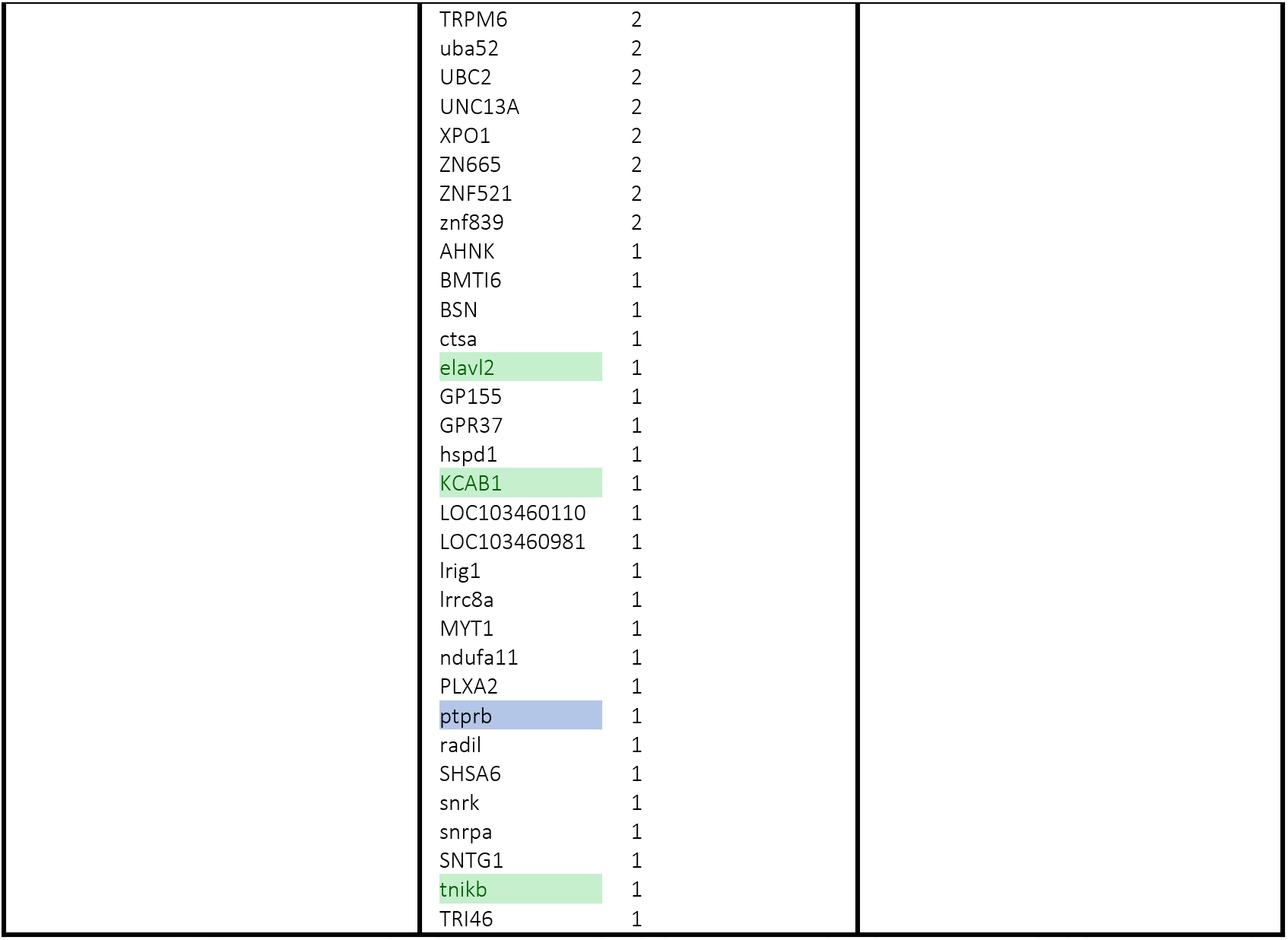
List of differentially coexpressed genes identified by BFDCA by brain region and line with number of differentially correlated connections (DC gene pairs) for each gene. Genes highlighted in green are DC in more than one comparison, and those highlighted in blue are also Diferentially Expressed.

**Table S9.**
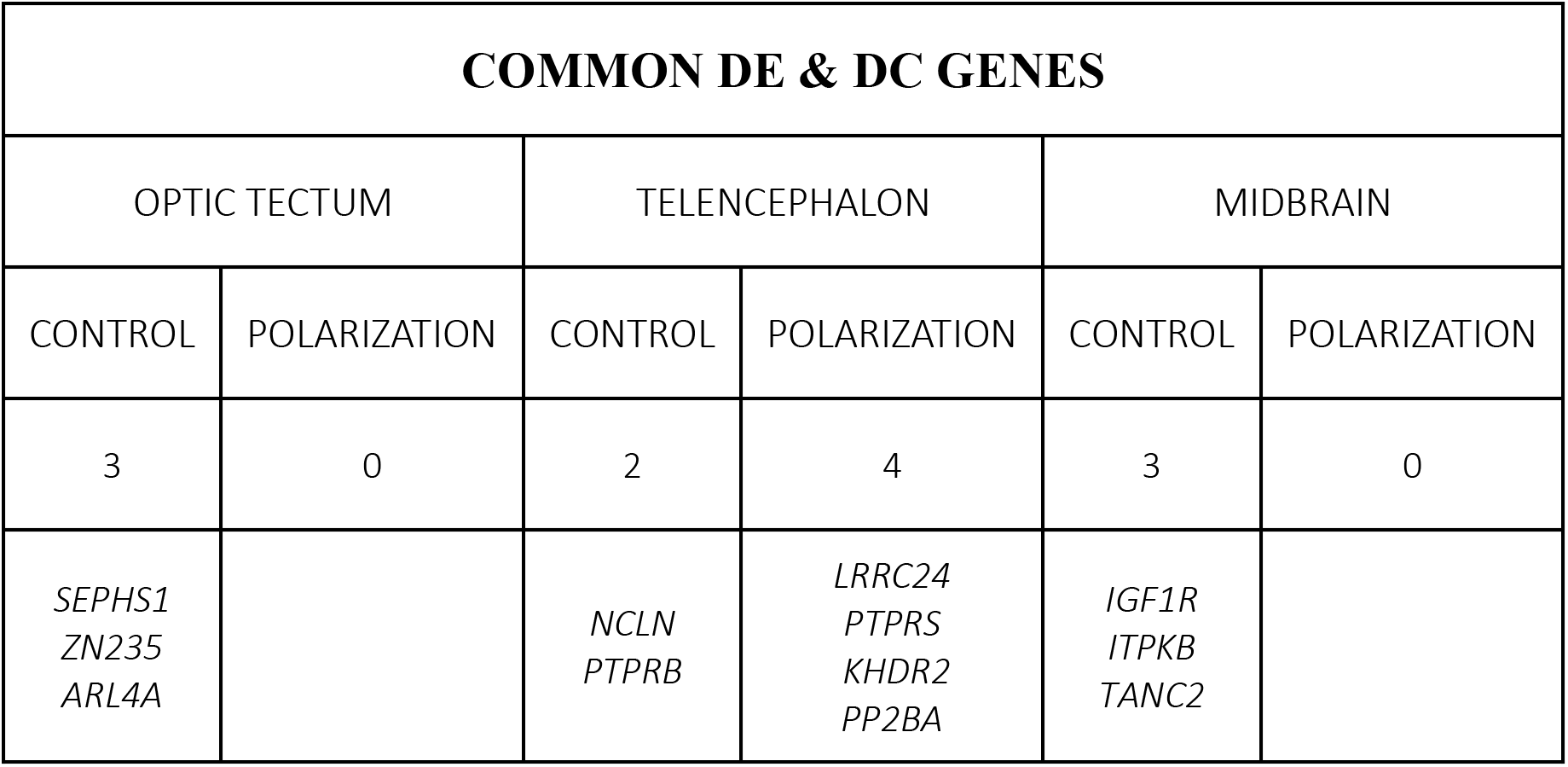
Summary of genes found to be differentially co-expressed and differentially expressed in our transcriptomic analyses

**Table S10.** KEGG pathways of interest associated in BFDCA differentially coexpressed gene pairs. Only showing pathways and select genes relevant to neural processes associated with behaviour that presented a FDR corrected p-value <0.05

| OPTIC TECTUM |  |
| --- | --- |
| Polarization-selection lines |  |
| Pathways of interest (enriched) | Genes of interest |
| Glutamatergic synapse | GLUL,GNG7,ITPR1 |
| Long-term depression | GRID2,ITPR1 |
| Dopaminergic synapse | GNG7,ITPR1 |
| GABAergic synapse | GLUL,GNG7 |
| Cholinergic synapse | GNG7,ITPR1 |
| Serotonergic synapse | GNG7,ITPR1 |
| Control lines |  |
| Signal Transduction of S1P Receptor | MAPK6 |
| TELENCEPHALON |  |
| Polarization-selection lines |  |
| Pathways of interest (enriched) | Genes of interest |
| FoxO signaling pathway | GRM1,MAPK9 |
| Phospholipase D signaling pathway | GRM1,RHEB |
| Synaptic vesicle cycle | SLC17A6,RAB3A |
| Glutamatergic synapse | SLC17A6,GRM1 |
| JNK (c-Jun kinases) phosphorylation and activation mediated by activated human TAK1 | MAPK9 |
| GABA synthesis, release, reuptake and degradation | RAB3A |
| Glutamate Neurotransmitter Release Cycle | RAB3A |
| Control lines |  |
| Glutamatergic synapse | CACNA1A,GRIA2,SLC1A1,GNAQ,GRM5,PRKCA,SLC1A2 |
| Synaptic vesicle cycle | CACNA1A,SLC1A1,SLC1A2,SLC6A11,UNC13A |
| PI3K-Akt signaling pathway | PKN1,ITGB5,BDNF,MAGI2,NTRK2,PHLPP2,PPP2R5C,PRKCA |
| Long-term depression | CACNA1A,GRIA2,GNAQ,PRKCA |
| Dopaminergic synapse | CACNA1A,GRIA2,GNAQ,PPP2R5C,PRKCA |
| Long-term potentiation | GRIA2,GNAQ,GRM5,PRKCA |
| Cholinergic synapse | ACHE,CACNA1A,GNAQ,PRKCA |
| Ras signaling pathway | HTR7,BDNF,NTRK2,RAC3,PRKCA |
| MIDBRAIN |  |
| Polarization-selection lines |  |
| EGFR tyrosine kinase inhibitor resistance | PDGFRB,IGF1R,NRG2 |
| Ras signaling pathway | PDGFRB,IGF1R,PLA1A |
| MAPK signaling pathway | PDGFRB,IGF1R,IL1A |
| AMPK signaling pathway | IGF1R,PFKFB2 |
| Control lines |  |
| cAMP signaling pathway | PPARA,GRIA2,ATP2B2,CNGB1 |
| Rap1 signaling pathway | MAGI1,IGF1R,RASSF5 |
| Long-term depression | IGF1R,GRIA2 |
| AMPK signaling pathway | IGF1R,PFKP |
| cGMP-PKG signaling pathway | ATP2B2,CNGB1 |
| Glutamate binding, activation of AMPA receptors and synaptic plasticity | GRIA2,GRIP1,TSPAN7 |

**Table S10:**
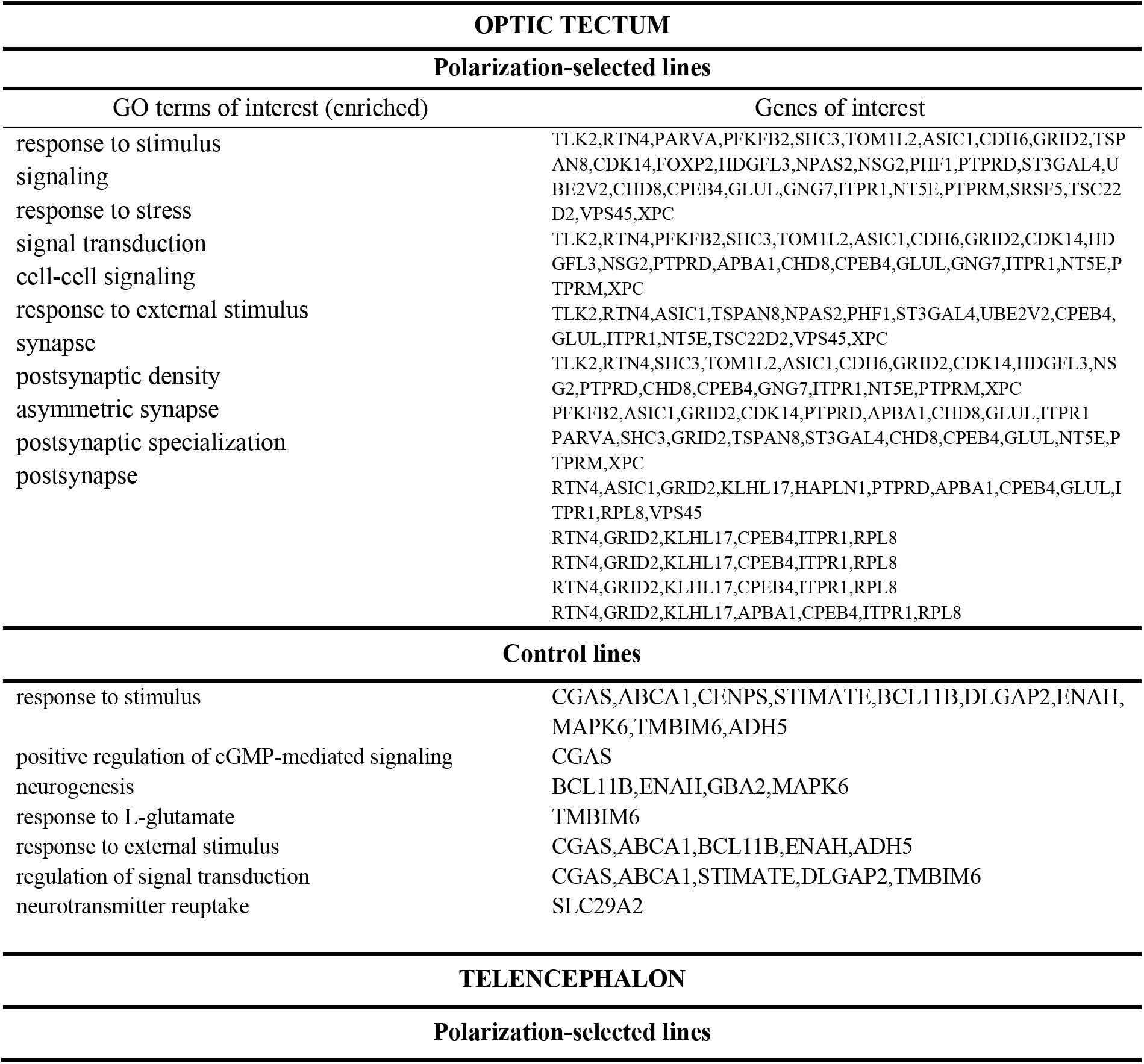
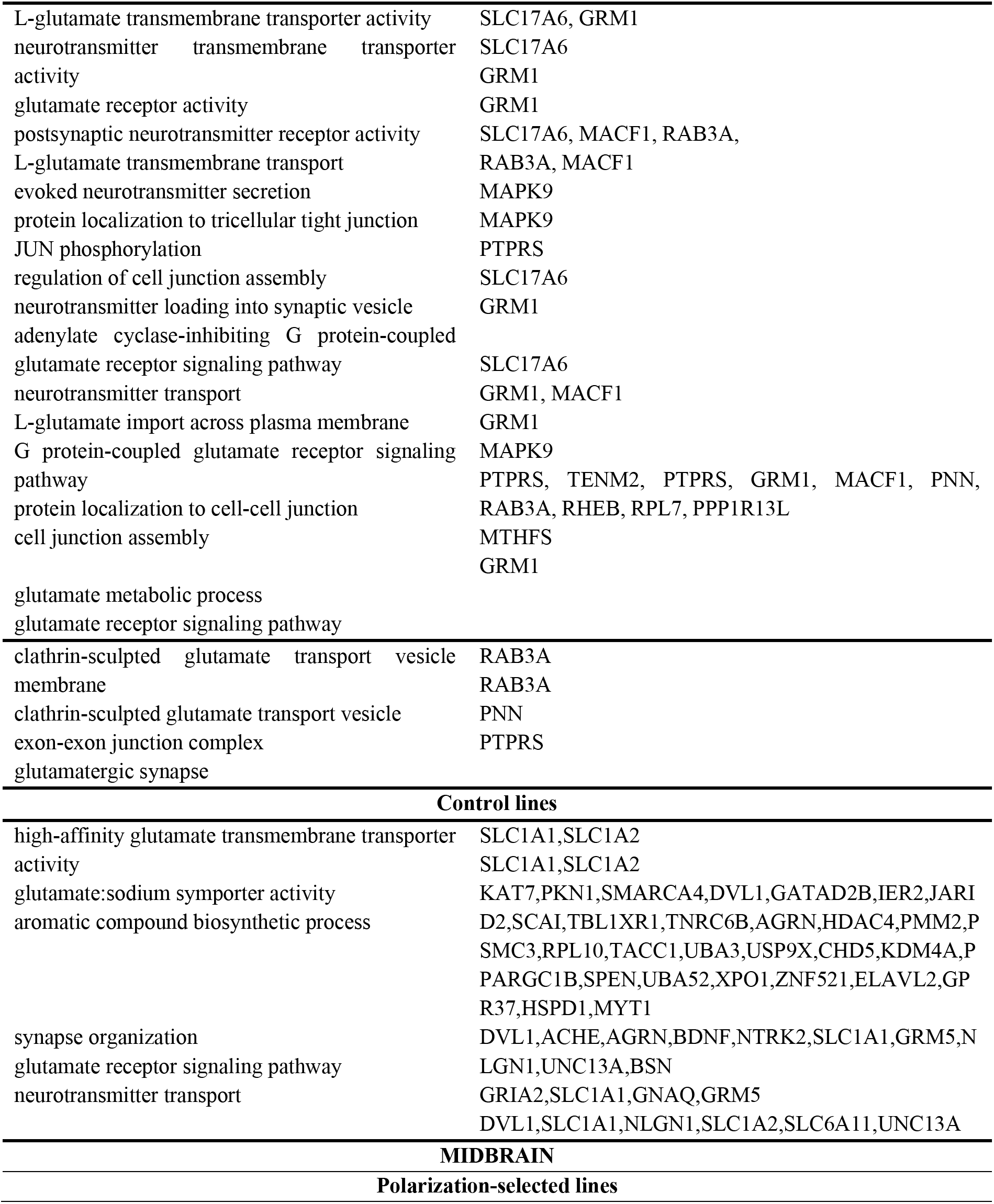
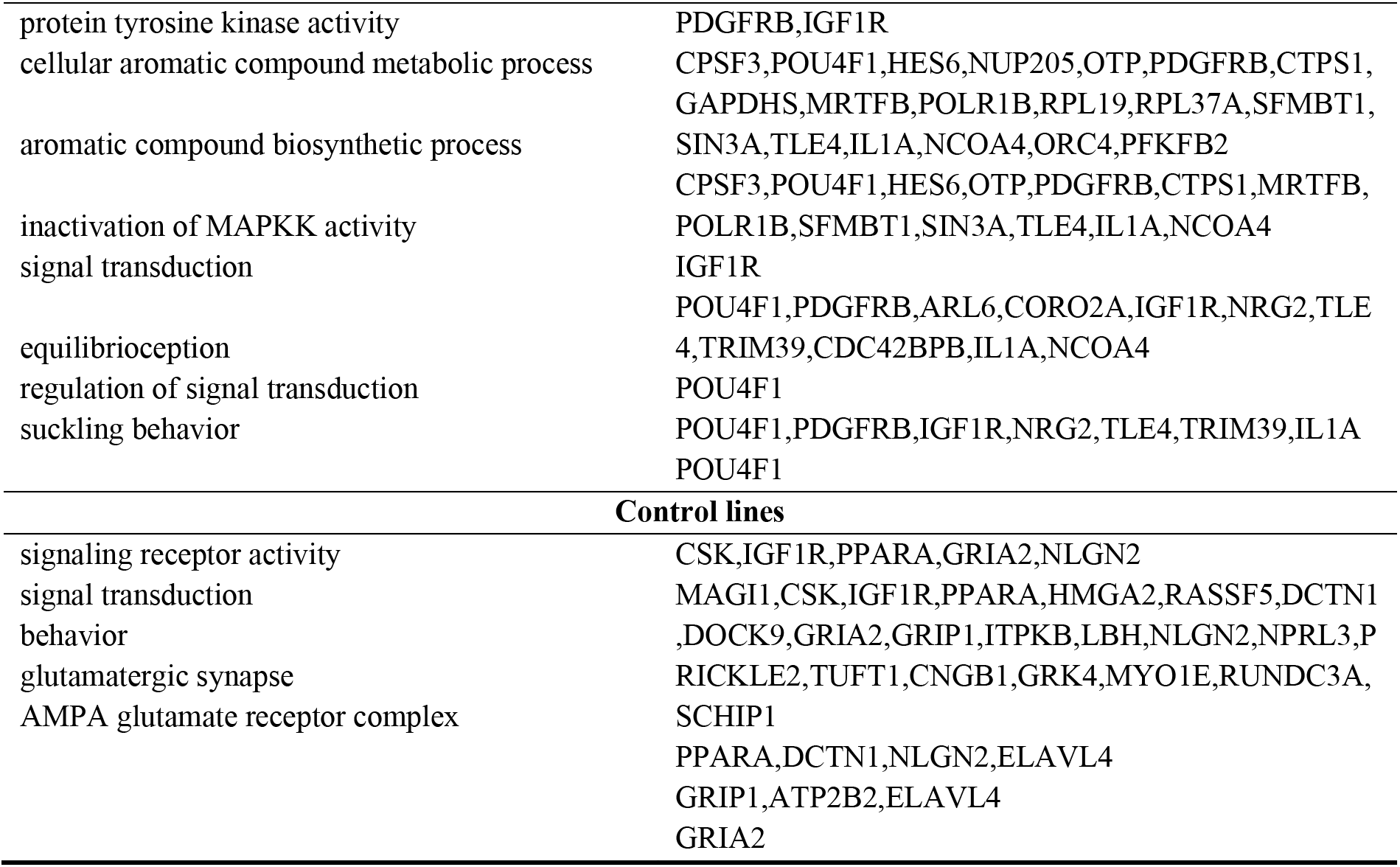
Gene Ontology (GO) term of interest associated BFDCA DC gene pairs

